# Bazedoxifene rescues sexually dimorphic autistic-like abnormalities in mice carrying a biallelic *MDGA1* mutation

**DOI:** 10.1101/2025.06.01.657278

**Authors:** Seungjoon Kim, Jinhu Kim, Byeong Chan Kim, Hyeonho Kim, Yeji Yang, Hea Ji Lee, Jin Young Kim, Ji Won Um, Jaewon Ko

## Abstract

In the accompanying study, we describe two pairs of *MDGA1* missense mutations (Val116Met/Ala688Val and Tyr635Cys/Glu756Gln) from two patients with autism spectrum disorders (ASDs) and how these mutations exert distinct abnormalities in biological processes during central nervous system development. Here, we generated knockin (KI) mice harboring the murine version of Tyr635Cys/Glu756Gln MDGA1 (*Mdga1*^Y636C/E751Q^) and performed extensive behavioral analyses. Male KI mouse pups and adults exhibited impaired ultrasonic vocalizations and sensorimotor gating (ASD-relevant behavioral deficits), and thus exhibited phenotypes different from those of male *Mdga1* conditional knockout (cKO) mice. In contrast, adult female KI mice did not exhibit a range of ASD-like behavioral abnormalities. Electrophysiological analyses performed using both juvenile and adult *Mdga1*^Y636C/E751Q^ mice revealed sexually dimorphic and developmental stage-dependent compromises of GABAergic synaptic inhibition. In addition, proteomics analyses showed that the phospho-proteomic phenotypes differed between *Mdga1*^Y636C/E751Q^ and *Mdga1*-cKO mice. Treatment of male *Mdga1*^Y636C/E751Q^ KI mice with the FDA-approved estrogen receptor modulator, bazedoxifene, restores the KI-related molecular, GABAergic synapse, and behavioral changes. Collectively, our results suggest a novel treatment strategy for ASDs that present developmentally regulated and/or sexually dimorphic features.

## Introduction

Synapse organization, which encompasses the formation, maintenance and elimination of synapses, is orchestrated by a plethora of various classes of synaptic proteins (1, 2). Among them, synaptic cell-adhesion molecules (CAMs) shape various forms of *trans*-cellular signaling pathways, coordinate the propagation of *trans*-synaptic signals in both presynaptic and postsynaptic neurons for proper synaptic assembly, and function as key lynchpins in dictating specific synaptic properties in the context of a neural circuit. Various candidate synaptic CAMs with putative synaptic functions have been identified and characterized for their abilities to promote various aspects of synapse development (2, 3). Intriguingly, certain synaptic CAMs are specifically expressed in either glutamatergic or GABAergic synapses, where they contribute to the construction of synapse type-specific signaling platforms. A subset of these synaptic CAMs is involved in regulating neural circuit-specific synaptic properties, either in line with their specific localization patterns (e.g., latrophilins and teneurin-3) or regardless of their distribution profiles (e.g., LRRTM3) (4-7). In contrast to the body of literature exploring the abilities of synaptic CAMs to facilitate synapse organization, only a few CAM-related mechanisms have been reported to negatively tune *trans*-synaptic signaling pathways (8).

The two members of the <u>m</u>eprin, A-5 protein, and receptor protein tyrosine phosphatase mu <u>d</u>omain-containing <u>g</u>lycosylphosphatidylinositol <u>a</u>nchor protein (MDGA) family proteins (MDGA1 and MDGA2) are highly expressed in the central nervous system (CNS) and presumed to be critical for early processes in the CNS development (8, 9). Remarkably, both MDGA paralogs suppress synapse maintenance in the adult CNS, as shown by analyses using transgenic mice lacking MDGA1 or MDGA2 (10). Both MDGA paralogs bind to members of the neuroligin (Nlgn) family of synaptic CAMs in *cis*-configuration and compete with presynaptic neurexins for binding to Nlgns (11-14).

However, MDGA1 targets amyloid precursor protein (APP), but not Nlgn2 (a Nlgn paralog exclusively expressed in GABAergic synapses) to negatively modulate GABAergic synapses in the hippocampal CA1 region (15). A recent systematic molecular replacement experiment performed using cultured hippocampal neurons from conditional knockout (cKO) mice lacking MDGA1 and/or MDGA2 demonstrated that MDGA1 and MDGA2 require completely different domains to execute their suppressive actions at GABAergic and glutamatergic synapses, respectively (10). MDGA1 requires its MAM domain to suppress synapse number, basal synaptic transmission, and synaptic strength at GABAergic synapses, whereas MDGA2 requires both Nlgn-binding activity and its MAM domain to regulate distinct excitatory synaptic properties. Intriguingly, conditional deletion of MDGA1 enhances synaptic strength but impairs GABA release probability in the dendritic, but not somatic, compartment of hippocampal CA1 pyramidal neurons *in vivo* (15). In addition, MDGA1-mediated suppression of GABAergic synapses depends on the presence of endogenous APP and is crucial for mediating proper novel object recognition memory in adult mice (15). Although MDGA1 has been reported to be linked to schizophrenia and bipolar disorder (16, 17), it remains unknown whether MDGA1 dysfunctions are indeed responsible for eliciting these neurodevelopmental disorders and, more importantly, which functions of MDGA1 are dysregulated in this context.

In the accompanying study, we identified two pairs of *MDGA1* missense mutations from patients with ASDs and found that these ASD-associated MDGA1 mutations induce different forms of impairment in CNS developmental processes. In the current study, we genetically introduced one of these mutations into mice (Tyr636Cys/Glu751Gln [Y636C/E751Q], corresponding to human Tyr635Cys/Glu756/Gln) and performed extensive behavioral, electrophysiological, and anatomical analyses. Our results showed that this knockin (KI) partly recapitulates a subset of *Mdga1* KO phenotypes, including abnormalities in ultrasonic vocalization and loss of GABAergic synaptic inhibition, but also yields distinct phenotypes. These phenotypes were sexually dimorphic and developmental stage-dependent related and could be at least partly rescued by the repurposed FDA-approved drug, bazedoxifene. Together, our findings highlight that improper synaptic inhibition appears to be a mechanism underlying ASDs, and suggest a novel potential treatment strategy for social communication deficits in ASDs.

## Results

### Adult male *Mdga1*^Y636C/E751Q^ KI mice exhibit different autistic-like behavioral abnormalities than adult male *Mdga1*-cKO mice

Our functional analyses revealed that the two ASD-associated MDGA1 substitutions induced abnormalities in distinct facets of neuronal and synaptic developmental processes (see the accompanying paper). However, *in utero* overexpression of MDGA1 Y635C/E756Q did not significantly impair pup isolation-induced USVs or affect neuronal migration (see the accompanying paper). To further investigate whether this substitution could influence behaviors in adult mice (**Fig. 1A**), we used a CRISPR/Cas9 system to generate *Mdga1* KI mice carrying Y636C and E751Q (Tyr636 and Glu751 in mouse *Mdga1* correspond to Tyr635 and Glu756 in human MDGA1) (see **Fig. 3A** for details). Our behavioral analyses showed that heterozygous *Mdga1*^Y636C/E751Q^ pups exhibited decreased USVs throughout development, similar to *Mdga1*-cKO pups (**Fig. 1B** and **1C**). In addition, homozygous *Mdga1*^Y636C/E751Q^ pups showed more severe impairment of USVs than their heterozygous counterparts (**Fig. 1B** and **1C**). These results imply that the presence of endogenous MDGA1 is likely to compensate for behavioral impairment under *in utero* electroporation of a vector encoding *Mdga1*^Y636C/E751Q^_._

**Figure 1.**
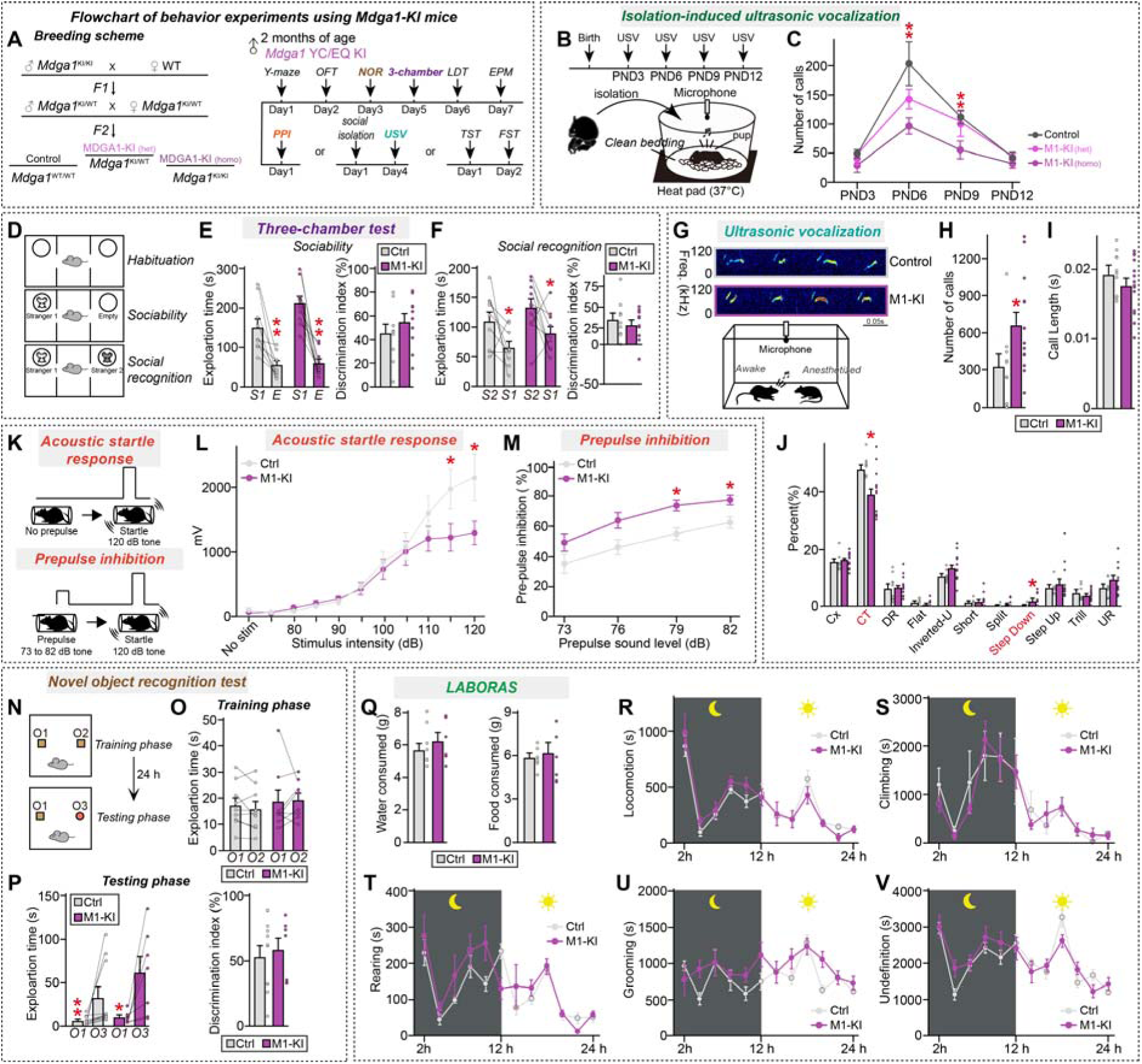
Analysis of mouse behavior in adult male *Mdga1*^Y636C/E751Q^ KI mice. (**A**) Breeding scheme followed to generate *Mdga1*^Y636C/E751Q^ KI mice, as well as the timeline of behavioral experiments performed on male MDGA1 KI mice, including Y-maze, open field test (OFT), novel object recognition (NOR), three-chamber test, light-dark transition (LDT), elevated plus maze (EPM), tail suspension test (TST), forced swim test (FST), social isolation, USV, and prepulse inhibition (PPI) measurements. (**B**) Experimental setup for recording USVs from isolated pups. Pups were separated from their mother and placed on clean bedding with a heat pad (37°C). A microphone was used to record the USVs. (**C**) Number of USV calls recorded from control and *Mdga1*^Y636C/E751Q^ KI pups at PND3, PND6, PND9 and PND12. Data are presented as means ± SEMs (n = 9–13 pups/group; \*\**p* < 0.01; Mann–Whitney *U* test). (**D–F**) Schematic (**D**) of the three-chamber social interaction test used to assess sociability and social recognition in mice. Quantification of exploration time during the three-chamber test for sociability (**E**; stranger 1 vs. empty) and social recognition (**F**; stranger 1 vs. stranger 2) in adult male control and *Mdga1*^Y636C/E751Q^ KI mice. Data are presented as means ± SEMs (n = 9–10 mice/group; \**p* < 0.05, \*\**p* < 0.01; Wilcoxon matched-pairs signed rank test). (**G–J**) Representative USV traces and schematic (**G**) of the recording setup for adult mice. Mice were placed in isolation, and USVs were recorded using a microphone. Number of USV calls (**H**) and call length (**I**) recorded from adult male control and *Mdga1*^Y636C/E751Q^ KI mice. Data are presented as means ± SEMs (n = 10–14 mice/group). Percent distribution of different USV call types (**J**; complex trill, downward ramp, flat, inverted-U, short, split, step down, step up, trill, upward ramp) in adult control and *Mdga1*^Y636C/E751Q^ KI mice. Data are presented as means ± SEMs (n = 10–14 mice/group; \**p* < 0.05; Mann–Whitney *U* test). (**K–M**) Schematic (**K**) of the PPI test. Mice were exposed to a prepulse sound (73 to 82 dB) followed by a startle pulse (120 dB), and the inhibition of the startle response was measured. (**L**) Acoustic startle response (**L**) in adult male control and *Mdga1*^Y636C/E751Q^ KI mice. Prepulse inhibition of the acoustic startle response (**M**) in adult male control and *Mdga1*^Y636C/E751Q^ KI mice. Data are presented as means ± SEMs (n = 12–13 mice/group; \**p* < 0.05; Mann–Whitney *U* test). (**N–P**) Schematic (**N**) of the NOR test. Training phase and testing phase timelines are shown, with object placements (O1, O2, O3) indicated. Quantification of exploration time during the training phase (**O**) and testing phase and the discrimination index (**P**) of the NOR test in control and *Mdga1*^Y636C/E751Q^ KI mice. Data are presented as means ± SEMs (n = 7–10 mice/group; \**p* < 0.05, \*\**p* < 0.01; Wilcoxon matched-pairs signed rank test). (**Q–V**) Quantification of various homecage activities in LABORAS cages, where mouse movements were monitored for 24 h. Shown are water and food consumption in grams (**Q**), and locomotion (**R**), climbing (**S**), rearing (**T**), grooming (**U**), and undefined (**V**) behavior of adult male control and *Mdga1*^Y636C/E751Q^ KI mice. Data were plotted every 2 h. The light and dark cycles of the day are represented by sun and moon icons, respectively. Data are presented as means ± SEMs (n = 6 mice/group).

When we subjected adult male *Mdga1*^Y636C/E751Q^ KI mice to a battery of behavioral tests similar to those used for *Mdga1*-cKO mice (**Fig. 1A**; see also accompanying paper), we found that both male adult *Mdga1*^Y636C/E751Q^ KI and *Mdga1*-cKO mice were viable and fertile, exhibiting no obvious abnormality, morbidity, or premature mortality (**Supplemental Fig. 1**). Male *Mdga1*^Y636C/751Q^ KI mice showed normal anxiety/exploration-related behavior, locomotor activity, working memory, and repetitive/compulsive-like behavior (**Supplemental Fig. 2**). Intriguingly, male *Mdga1*^Y636C/751Q^ KI mice exhibited a decreased acoustic startle response (ASR) but increased prepulse inhibition (PPI) (**Fig. 1K– 1M**), and thereby differed from the male *Mdga1*-cKO mice (see accompanying paper). Male *Mdga1*^Y636C/751Q^ KI mice exhibited normal sociability and object recognition memory (**Fig. 1D–1F** and **Fig. 1N–1P**) but, similar to their male cKO counterparts, male *Mdga1*^Y636C/751Q^ KI mice encountering female mice exhibited increased USV calls and these USVs had an altered syllable composition (**Fig. 1G–1J**). Overall, male *Mdga1*^Y636C/751Q^ KI mice exhibited a narrower range of behavioral impairments than male *Mdga1*-cKO mice, particularly in sensorimotor gating and sociability. Again, the female *Mdga1*^Y636C/751Q^ KI mice failed to exhibit any of the behavioral deficits seen in their male KI counterparts, further highlighting the sexually dimorphic manifestation of MDGA1 dysfunctions (**Fig. 2**).

**Figure 2.**
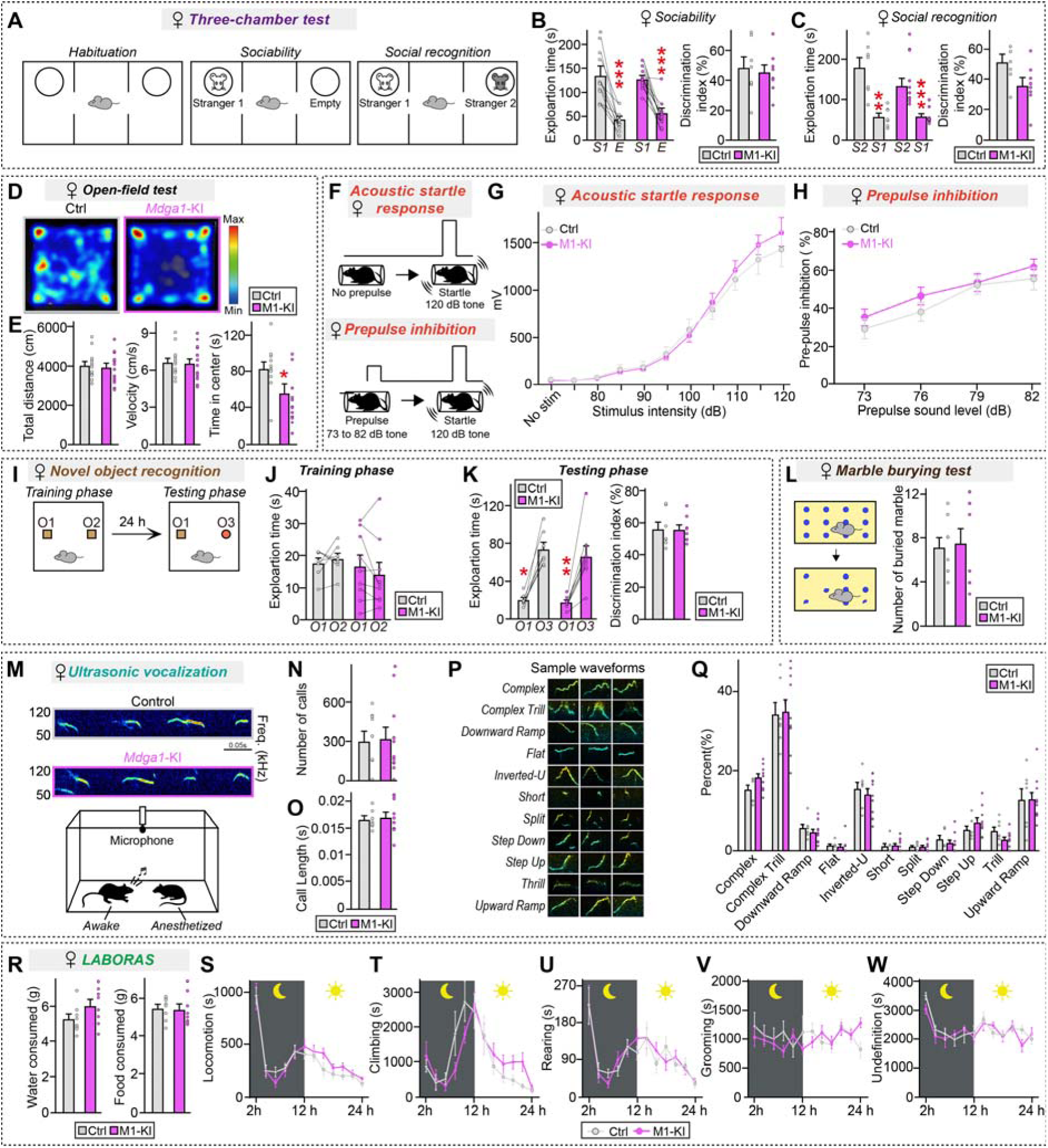
Analysis of behaviors of adult female *Mdga1*^Y636C/E751Q^ KI mice. (**A–C**) Three-chamber test results showing a schematic of the test (**A**) and the exploration time and discrimination index during the sociability (**B**; stranger 1 vs. empty) and social recognition (**C**; stranger 1 vs. stranger 2) phases for female control and *Mdga1*^Y636C/E751Q^ KI mice. Data are presented as means ± SEMs (n =11–13 mice/group; \*\**p* < 0.01, \*\*\**p* < 0.001; Wilcoxon matched-pairs signed rank test). (**D** and **E**) Open-field test results showing total distance traveled, velocity, and time spent in the center for male control and *Mdga1*^Y636C/E751Q^ KI mice. Data are presented as means ± SEMs (n = 11–13 mice/group; \**p* < 0.05, Mann–Whitney *U* test). (**F–H**) Schematic (**F**) of the prepulse inhibition (PPI) test. Mice were exposed to a prepulse sound (73 to 82 dB) followed by a startle pulse (120 dB), and the inhibition of the startle response was measured.) Acoustic startle response (**G**) in female control and male *Mdga1*^Y636C/E751Q^ KI mice. Data are presented as means ± SEMs (n = 11–14 mice/group). PPI of the acoustic startle response (**H**) in female control and MDGA1 KI mice. Data are presented as means ± SEMs (n = 11–14 mice/group). (**I–K**) Novel object recognition test results showing a schematic of the test (**I**) and the exploration time and discrimination index during the training (**J**) and testing phases (**K**) for male control and *Mdga1*^Y636C/E751Q^ KI mice. Data are presented as means ± SEMs (n = 7–9 mice/group; \**p* < 0.05, \*\**p* < 0.01; Wilcoxon matched-pairs signed rank test). (**L**) Marble burying test results showing the number of marbles buried by female control and female *Mdga1*^Y636C/E751Q^ KI mice. Data are presented as means ± SEMs (n = 8 mice/group). (**M–Q**) Ultrasonic vocalization results showing a schematic of the test (**M**), the number of calls (**N**) and the call length (**O**) for male control and *Mdga1*^Y636C/E751Q^ KI mice. Representative waveforms (**P**) and distribution of different call types (**Q**) are presented. Data are presented as means ± SEMs (n = 16– 18 mice/group). (**R–W**) LABORAS test results showing various behaviors: water and food consumption (**R**), locomotion (**S**), climbing (**T**), rearing (**U**), grooming (**V**), and undefined activities (**W**) for male control and *Mdga1*^Y636C/E751Q^ KI mice measured over 24 hours. Data are presented as means ± SEMs (n = 8 mice/group).

**Figure 3.**
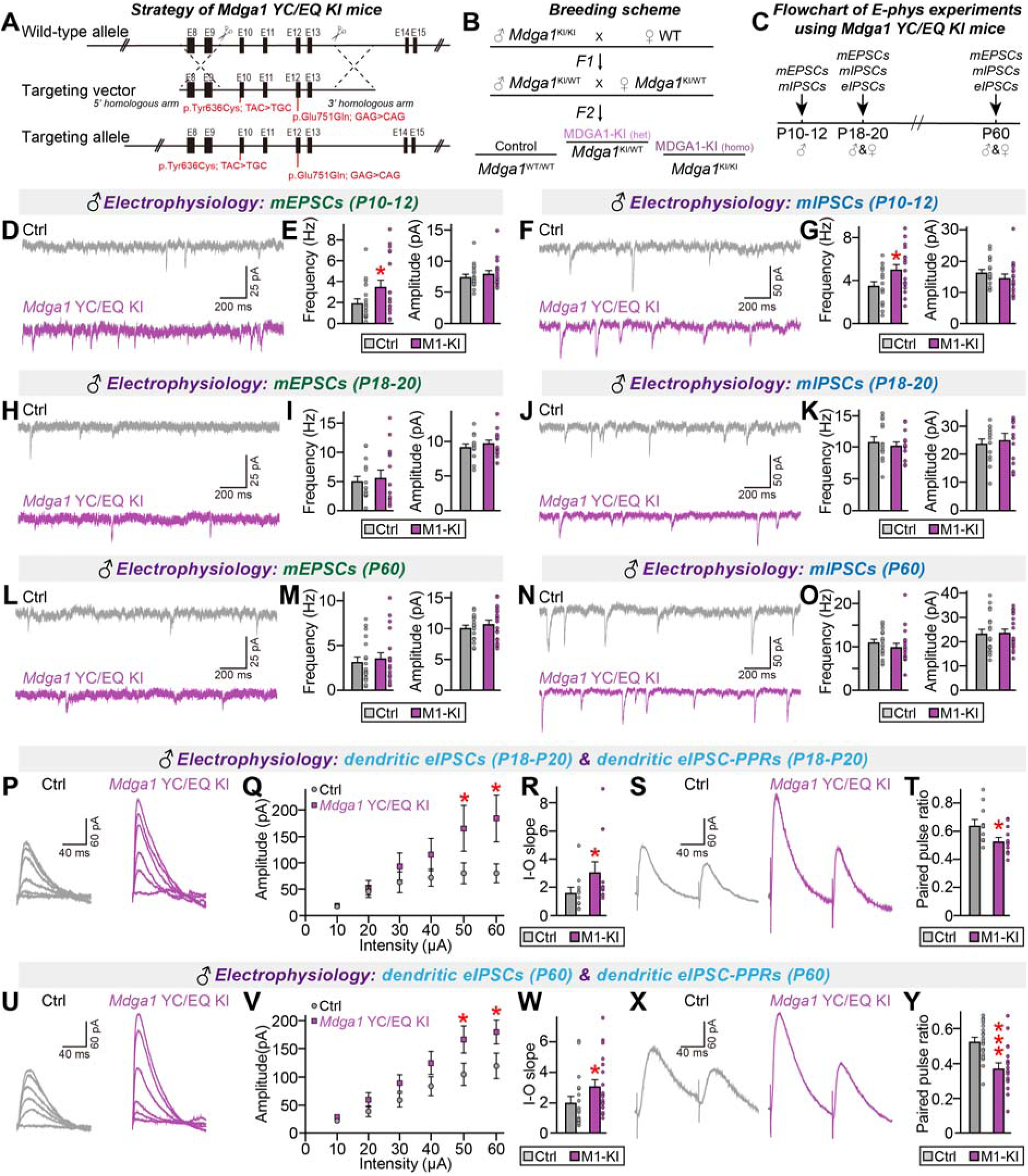
Analysis of GABAergic synaptic properties in the hippocampal CA1 pyramidal neurons of *Mdga1*^Y636C/E751Q^ KI mice. (**A**) Schematic illustrating the genetic targeting strategy used to generate *Mdga1*^Y636C/E751Q^ KI mice. The point mutations p.Tyr636Cys (TAC>TGC) and p.Glu751Gln (GAG>CAG) are shown on the targeting allele and WT allele. (**B**) Breeding scheme followed to generate *Mdga1*^Y636C/E751Q^ KI mice. (**C**) Flowchart of the electrophysiology experiments conducted on *Mdga1*^Y636C/E751Q^ KI mice at different developmental stages (P10–12, P18–20 and P60). (**D** and **E**) Representative traces (**D**) and quantification (**E**) of mEPSCs in CA1 pyramidal neurons from control and *Mdga1*^Y636C/E751Q^ KI mice at P10–12. Data are presented as means ± SEMs (n = 19 cells/group; \**p* < 0.05; Mann–Whitney *U* test). (**F** and **G**) Representative traces (**F**) and quantification (**G**) of mIPSCs in CA1 pyramidal neurons from control and *Mdga1*^Y636C/E751Q^ KI mice at P10–12. Data are presented as means ± SEMs (n = 18–19 cells/group; \**p* < 0.05; Mann–Whitney *U* test). (**H** and **I**) Representative traces (**H**) and quantification (**I**) of mEPSCs in CA1 pyramidal neurons from control and *Mdga1*^Y636C/E751Q^ KI mice at P18–20. Data are presented as means ± SEMs (n = 15 cells/group). (**J** and **K**) Representative traces (**J**) and quantification (**K**) of mIPSCs in CA1 pyramidal neurons from control and *Mdga1*^Y636C/E751Q^ KI mice at P18–20. Data are presented as means ± SEMs (n = 14 cells/group). (**L** and **M**) Representative traces (**L**) and quantification (**M**) of mEPSCs in CA1 pyramidal neurons from control and *Mdga1*^Y636C/E751Q^ KI mice at P60. Data are presented as means ± SEMs (n = 19 cells/group). (**N** and **O**) Representative traces (**N**) and quantification (**O**) of mIPSCs in CA1 pyramidal neurons from control and *Mdga1*^Y636C/E751Q^ KI mice at P60. Data are presented as means ± SEMs (n = 19 cells/group). (**P–T**) Representative traces (**P** and **S**) and quantification of dendritic eIPSCs in CA1 pyramidal neurons from control and *Mdga1*^Y636C/E751Q^ KI mice at P18–20. Input-output (I-O) curves (**Q** and **R**) and paired-pulse ratio (PPR) (**T**) of dendritic eIPSCs. Data are presented as means ± SEMs (n = 11–12 cells/group; \**p* < 0.05; Mann–Whitney *U* test). (**U–Y**) Representative traces (**U** and **X**) and quantification of dendritic evoked inhibitory postsynaptic currents (eIPSCs) in CA1 pyramidal neurons from control and *Mdga1*^Y636C/E751Q^ KI mice at P60. Input-output (I-O) curves (**V** and **W**) and paired-pulse ratio (PPR) (**Y**) of dendritic eIPSCs. Data are presented as means ± SEM (n = 19 cells/group; \**p* < 0.05; Mann–Whitney *U* test).

### Adult male *Mdga1*^Y636C/E751Q^ KI mice phenocopy the electrophysiological abnormalities of ***Mdga1*-cKO mice**

We additionally analyzed the levels of MDGA1 and other synaptic proteins in *Mdga1*^Y636C/E751Q^ KI and cKO mice. Quantitative RT-PCR analyses revealed that forebrain regions (hippocampus and cortex), but not cerebellum, exhibited marked decreases in the mRNA levels of *Mdga1* with no change in *Mdga2* (**Supplemental Fig. 3A**). Semi-quantitative immunoblotting experiments showed that *Mdga1*-cKO caused a significant loss of MDGA1, whereas the Y636C/R751Q substitution did not affect the protein level of MDGA1 (**Supplemental Fig. 3B** and **3C**). We also observed that the expression levels of GABA_A_Rγ2 and vesicular GABA transporter (VGAT) were increased in cKO and KI mice, that of gephyrin was substantially increased in cKO mice, and that of Nlgn2 was increased in KI mice (**Supplemental Fig. 3B** and **3C**). We did not observe any significant change in the levels of the other examined proteins, including those characteristics of excitatory synapses. These data suggest that *Mdga1*^Y636C/E751Q^ KI and cKO do not cause global changes in the molecular composition of the brain, except for the small increases seen in inhibitory synaptic markers among both KI and cKO mice.

We further performed immunohistochemical analyses using antibodies against VGAT and gephyrin (to label inhibitory synapses) or VGLUT1 and PSD-95 (to label excitatory synapses) in the hippocampus and mPFC of adult cKO and KI mice. Increased density of VGAT^+^ puncta was specifically observed in the *stratum lacunosum moleculare* layer of the hippocampus in adult male cKO and KI mice (**Supplemental Figs. 4** and **5**), but not their female counterparts. No change was observed among inhibitory and excitatory synapses in mPFC layer II/III of adult cKO and KI mice.

**Figure 4.**
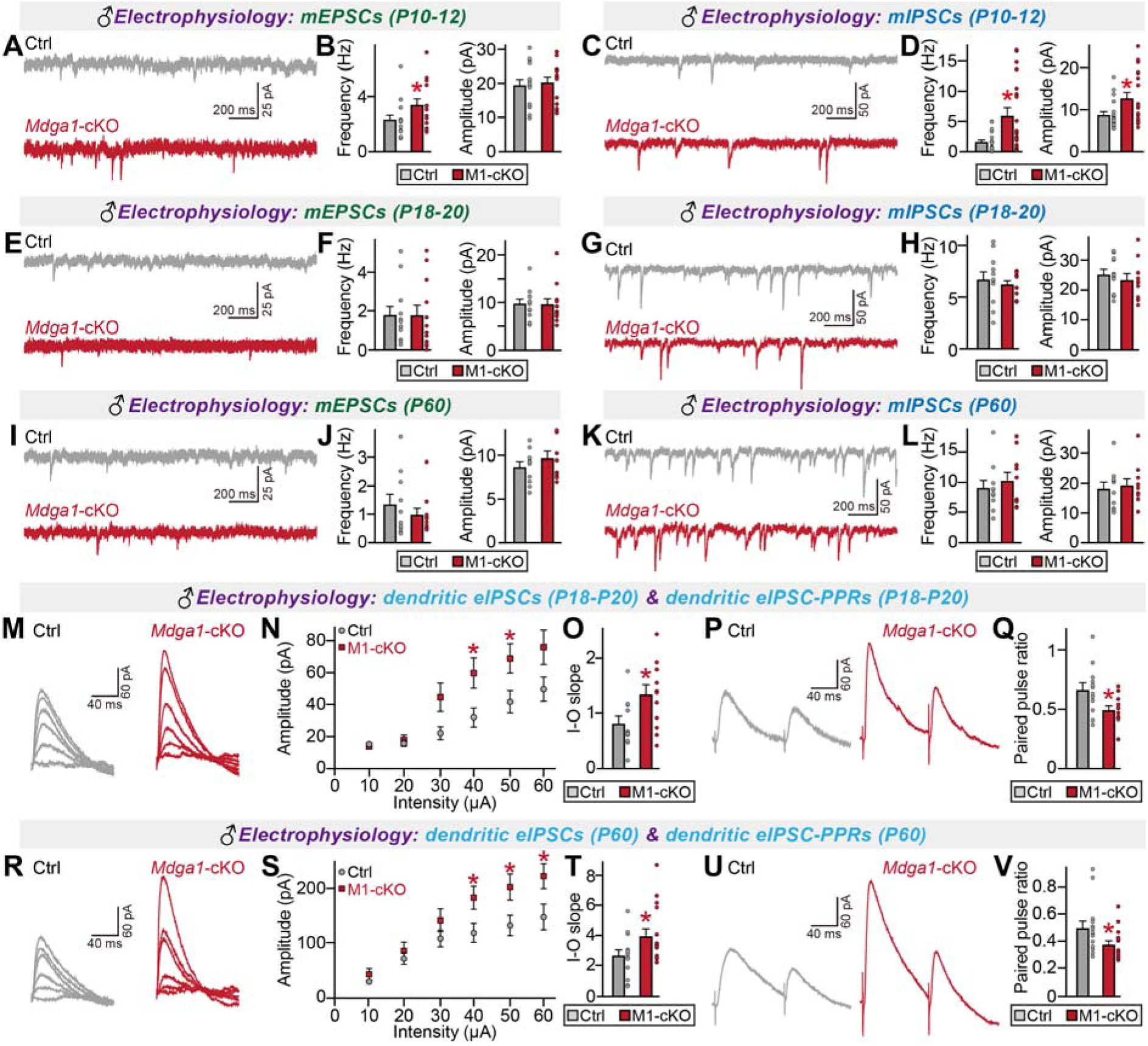
Analysis of electrophysiological properties in the hippocampal CA1 pyramidal neurons of various aged male *Mdga1*-cKO mice. (**A** and **B**) Representative traces (**A**) and averages of mEPSC frequencies and amplitudes from hippocampal CA1 pyramidal neurons of male control and *Mdga1*-cKO mice at P10–P12 (**B**; control, n = 15/6; M1-cKO, n = 14/6; where n denotes the number of cells/mice). Data are presented as means ± SEMs. (**C** and **D**) Representative traces (**C**) and averages of mIPSC frequencies and amplitudes from CA1 pyramidal neurons of male control and *Mdga1*-cKO mice at P10–P12 (**D**; control, n = 19/6; M1-cKO, n = 19/6). Data are presented as means ± SEMs. (**E** and **F**) Representative traces (**E**) and averages of mEPSC frequencies and amplitudes (**F**) from CA1 pyramidal neurons of male control and *Mdga1*-cKO mice at P18–P20 (control, n = 12/4; M1-cKO, n = 12/4). Data are presented as means ± SEMs. (**G** and **H**) Representative traces (**G**) and averages of mIPSC frequencies and amplitudes (**H**) from CA1 pyramidal neurons of male control and *Mdga1*-cKO mice at P18–P20 (control, n = 11/4; M1-cKO, n = 10/4). Data are presented as means ± SEMs. (**I** and **J**) Representative traces (**I**) and averages of mEPSC frequencies and amplitudes (**J**) from CA1 pyramidal neurons of male control and *Mdga1*-cKO mice at P60 (control, n = 10/4; M1-cKO, n= 10/4). Data are presented as means ± SEMs. (**K** and **L**) Representative traces (**K**) and averages of mIPSC frequencies and amplitudes (**L**) from CA1 pyramidal neurons of male control and *Mdga1*-cKO mice at P60 (control, n = 10/4; M1-cKO, n = 10/4). Data are presented as means ± SEMs. (**M–O**) Representative traces (**M**) and averages of dendritic eIPSCs (**N** and **O**) from CA1 pyramidal neurons of male control and *Mdga1*-cKO mice at P18–P20 (control, n = 12/4; M1-cKO, n = 12/4). Data are presented as means ± SEMs. (**P** and **Q**) Representative traces (**P**) and averages of dendritic eIPSC-PPRs (**Q**) from CA1 pyramidal neurons of male control and *Mdga1*-cKO mice at P18–P20 (control, n = 13/4; M1-cKO, n = 13/4). Data are presented as means ± SEMs. (**R–T**) Representative traces (**R**) and averages of dendritic eIPSCs (**S** and **T**) from CA1 pyramidal neurons of male control and *Mdga1*-cKO mice at P60 (control, n = 13/5; M1-cKO, n = 16/5). Data are presented as means ± SEMs. (**U** and **V**) Representative traces (**U**) and averages of dendritic eIPSC-PPRs (**V**) from CA1 pyramidal neurons of male control and *Mdga1*-cKO mice at P60 (control, n = 14/5; M1-cKO, n = 16/5). Data are presented as means ± SEMs (\**p* < 0.05; two-tailed non-parametric Mann–Whitney *U* test).

**Figure 5.**
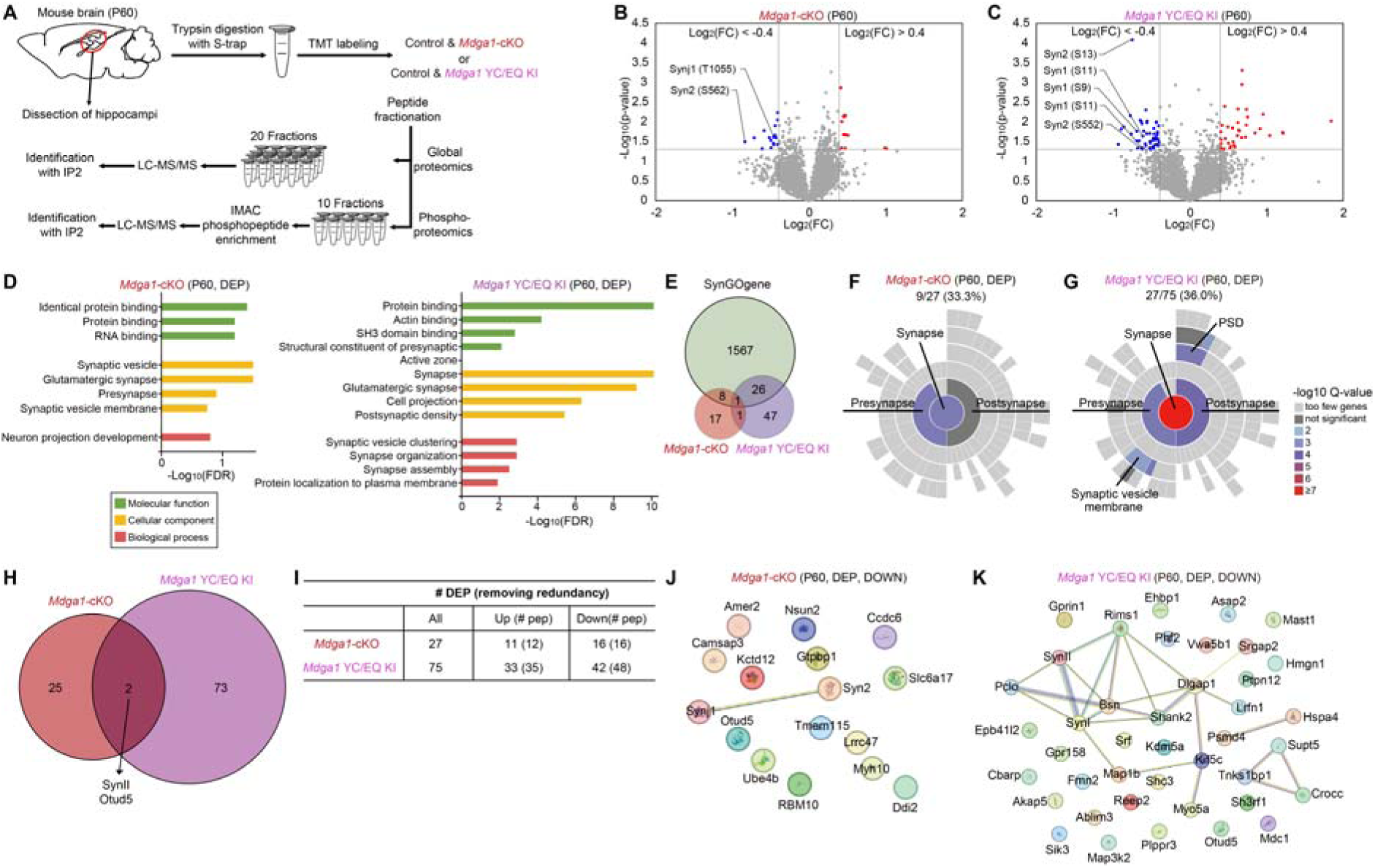
Phosphoproteomic analysis of hippocampi from adult *Mdga1*-cKO and Mdga*^1Y636C/E751Q^* KI mice. (A) Schematic diagram of phosphoproteomic analysis of hippocampal lysates from adult male *Mdga1*-cKO, *Mdga1*^Y636C/E751Q^ KI, and littermate control mice. (B) Volcano plot of 4,338 phosphopeptides identified from hippocampal lysates of adult male *Mdga1*-cKO mice. Differentially expressed phosphopeptides (DEPPs) with significant increases or decreases (Log_2_FC < -0.4 or > 0.4; *p* < 0.05) are shown in red (12 phosphopeptides) and blue (16 phosphopeptides), respectively. (C) Volcano plot of 5,124 phosphopeptides identified from hippocampal lysates of adult male *Mdga1*^Y636C/E751Q^ KI mice. DEPPs with significant increases or decreases (Log_2_FC < -0.4 or > 0.4; *p* < 0.05) are shown in red (35 phosphopeptides) and blue (48 phosphopeptides), respectively. (D) Gene ontology (GO) analysis was performed on differentially expressed phosphoproteins (DEPs) identified from the phosphoproteomic analyses of adult male *Mdga1*-cKO (**left**) and *Mdga1*^Y636C/E751Q^ KI (**right**) mice. The bar charts depict the significantly enriched GO terms of the Molecular Function (green), Cellular Component (yellow), and Biological Process (red) categories, represented as - log_10_(FDR) values. (E) Venn diagram visualization of SynGO category proteins. (F) Localization annotations of SynGO category proteins from adult male *Mdga1*-cKO mice. (G) Localization annotations of SynGO category proteins from adult male *Mdga1*^Y636C/E751Q^ KI mice. (H) Venn diagram showing the overlap of DEPs between *Mdga1*-cKO and *Mdga1*^Y636C/E751Q^ KI mice. Two proteins, SynII and Otud5, are shared between condition; 25 DEPs are unique to *Mdga1*-cKO mice and 73 are unique to *Mdga1*^Y636C/E751Q^ KI mice. (I) Summary table of DEPs after removal of redundancy. The table shows the total number DEPs with a breakdown of upregulated and downregulated proteins in *Mdga1*-cKO and *Mdga1*^Y636C/E751Q^ KI mice, together with the number of unique peptides (# pep). (J) STRING analysis of downregulated DEPs from *Mdga1*-cKO mice (*p* < 0.05 and Log_2_FC < -0.4). (K) STRING analysis of downregulated DEPs from *Mdga1*^Y636C/E751Q^ KI mice (*p* < 0.05 and Log_2_FC < -0.4).

Next, we examined whether the *Mdga1*^Y636C/E751Q^ variant retains the ability of endogenous MDGA1 to regulate synaptic transmission in hippocampal CA1 pyramidal neurons (**Fig. 3A–3C**). Male KI mice were subjected to electrophysiological recordings for measurement of spontaneous synaptic transmission, evoked synaptic strength, and neurotransmitter release probability. Since growth stage-related differences in electrophysiological phenotypes were previously reported for *Mdga1*-KO mice (15, 18), we examined both juvenile (further subdivided into postnatal [P] 10–12 or P18–20) and adult (P60) male *Mdga1*^Y636C/E751Q^ mice (**Fig. 3C**). Whole cell patch-clamp recordings revealed no marked alteration in the frequency/amplitude of mIPSCs or mEPSCs in hippocampal CA1 neurons from juvenile and adult male *Mdga1*^Y636C/E751Q^ mice, with the exception that P10–12 *Mdga1*^Y636C/E751Q^ neurons exhibited increased mIPSC and mEPSC frequencies (**Fig. 3D–3O**). Parallel recordings obtained from male *Mdga1*-cKO mice were aligned with those from same-stage male *Mdga1*^Y636C/E751Q^ mice, showing increased mIPSC and mEPSC frequencies from P10–12 CA1 pyramidal neurons, but no change in spontaneous synaptic transmission from P18–20 or P60 CA1 pyramidal neurons (**Fig. 4**).

Hippocampal CA1-specific *Mdga1*-cKO mice were reported to exhibit increased dendritic IPSC amplitudes with increased PPRs specifically upon stimulation of distal dendrites (15). We thus examined whether these electrophysiological features were replicated in male *Mdga1*^Y636C/E751Q^ CA1 pyramidal neurons. Both P18–20 and P60 CA1 pyramidal neurons from male *Mdga1*^Y636C/E751Q^ KI mice exhibited marked increases in eIPSC amplitudes and IPSC-PPRs (**Fig. 3P–3Y**), as did the corresponding neurons from male *Mdga1*-cKO mice (**Fig. 4**). These findings reinforce the idea that the Y636C/E751Q substitution leads to loss-of-function effects. Unexpectedly, we observed a significant increase in dendritic aIPSC amplitudes in CA1 pyramidal neurons from male *Mdga1*^Y636C/E751Q^ KI and *Mdga1*-cKO mice (**Supplemental Fig. 6A–6D**). However, measurement of somatic aIPSCs revealed a specific increase in male *Mdga1*^Y636C/E751Q^ KI, but not male *Mdga1*-cKO mice (**Supplemental Fig. 6E– 6H**). In females, CA1 pyramidal neurons from adult female *Mdga1*^Y636C/E751Q^ KI and *Mdga1*-cKO mice displayed no alterations in eIPSC amplitudes or IPSC-PPRs, compared to control neurons (**Supplemental Fig. 7**). However, CA1 neurons from juvenile female *Mdga1*^Y636C/E751Q^ KI and *Mdga1*-cKO mice showed increased eIPSC amplitudes and IPSC-PPRs (**Supplemental Fig. 7**). These results suggest that the sex-specific changes in GABAergic synaptic strength reflect developmental effects and are consistent with the lack of detectable impairment in certain social behaviors among adult female *Mdga1*^Y636C/E751Q^ KI and *Mdga1*-cKO mice.

**Figure 6.**
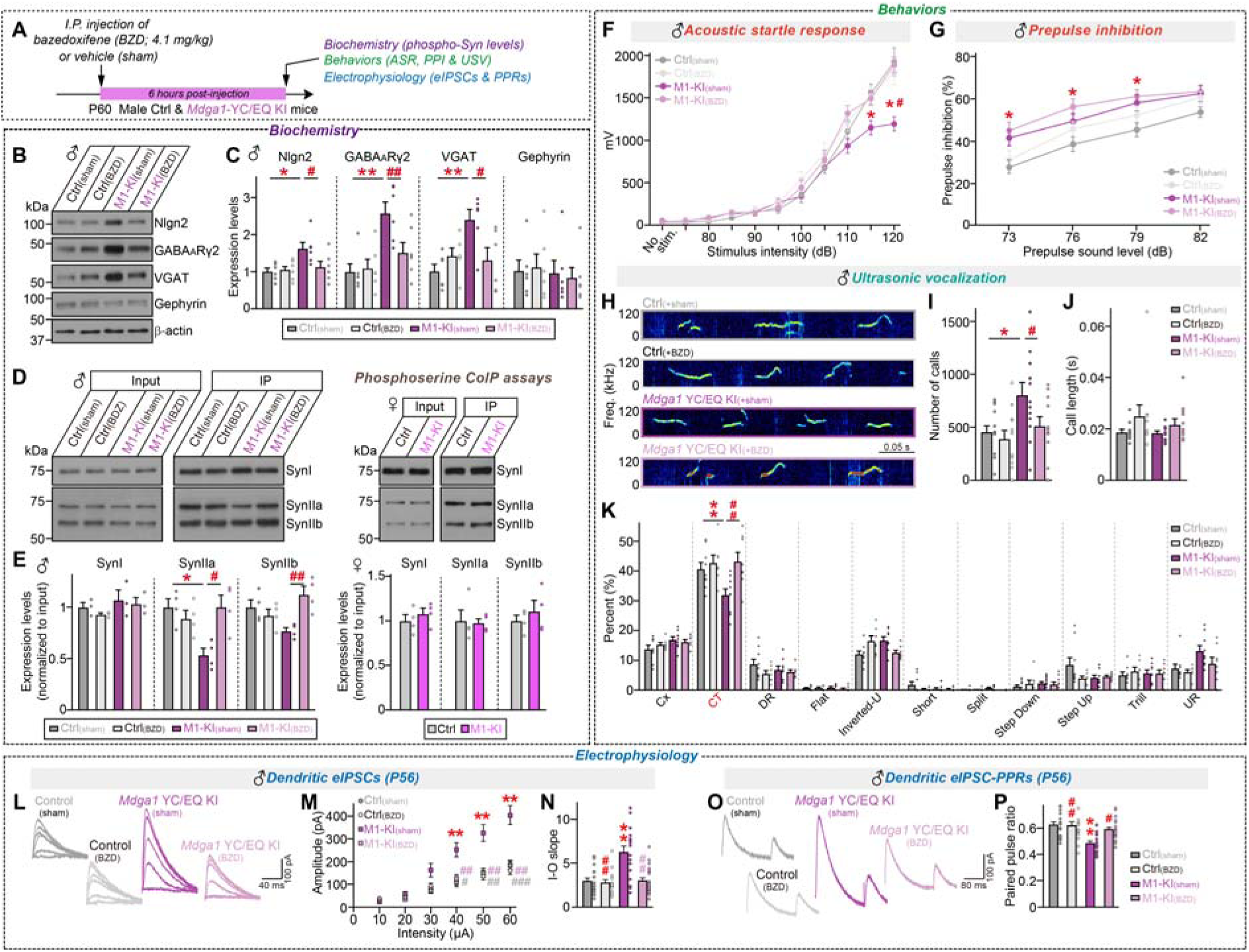
Bazedoxifene rescues sexually dimorphic alterations in the hippocampal CA1 pyramidal neurons of *Mdga1*^Y636C/E751Q^ KI mice. (A) Schematic of the experimental design for application of bazedoxifene (BZD) or vehicle (sham) to adult male control and *Mdga1*^Y636C/E751Q^ KI mice followed 6 hours later by various functional analyses. (B) Semi-quantitative immunoblotting of various synaptic proteins in the hippocampi of adult male control and *Mdga1*^Y636C/E751Q^ mice given BZD or sham treatment. β-actin was used as a loading control for normalization. (C) Semi-quantitative analysis of the immunoblotting results shown in (**B**). Expression levels of various proteins were normalized to that of β-actin and are presented as fold changes relative to those from control mice. Data are presented as means ± SEM (n = 6 per group; \**p* < 0.05, \*\**p* < 0.01, ^#^*p* < 0.05, ^##^*p* < 0.01; ‘#’ indicates statistical comparisons with their counterparts; two-way ANOVA followed by Tukey’s *post-hoc* test). (D) Representative immunoblot images showing phosphoprotein levels of synapsin I (SynI) or SynII from hippocampal lysates of adult male control and *Mdga1*^Y636C/E751Q^ KI mice given BZD or sham treatment. (E) Quantification of the SynI, SynIIa, and SynIIb immunoreactivities from anti-pSer immunoprecipitates, normalized to the total amount of the indicated Syn proteins in each group. Data are presented as means ± SEM (number of independent experiments; n = 4 for all experimental groups; *\*p* < 0.05, *^#^p* < 0.05, *^##^p* < 0.01; ‘#’ indicates statistical comparisons with their counterparts; two-way ANOVA followed by Tukey’s *post-hoc* test). (**F** and **G**) ASRs (**F**) and PPI of ASR (**G**) in adult male control and *Mdga1*^Y636C/E751Q^ mice given BDZ or sham treatment. Data are presented as means ± SEM (n = 20–21 mice/group; \**p* < 0.05, ^#^*p* < 0.05, ‘#’ indicates statistical comparisons with their counterparts; two-way ANOVA followed by Tukey’s *post-hoc* test). (**H–K**) For recordings obtained from adult male control and *Mdga1*^Y636C/E751Q^ KI mice given BZD or sham treatment: representative USV traces (**H**), numbers of USV calls (**I**), call lengths (**J**), and percent distribution of different USV call types (**K**). Data are presented as means ± SEM (n = 9–10 mice/group; \**p* < 0.05, \*\**p* < 0.01, ^##^*p* < 0.01, ‘#’ indicates statistical comparisons with their counterparts; two-way ANOVA followed by Tukeyʹs *post*-*hoc* test). (**L–P**) Representative traces (**L** and **O**) and quantification of dendritic eIPSCs in CA1 pyramidal neurons from control and *Mdga1*^Y636C/E751Q^ KI mice at P56. Input-output (I-O) curves (**M** and **N**) and PPR (**P**) of dendritic eIPSCs. Data are presented as means ± SEM (n = 14–21 cells/group; \*\**p* < 0.01, ^#^*p* < 0.05, ^##^*p* < 0.01; ‘#’ indicates statistical comparisons with their counterparts; two-way ANOVA followed by Tukey’s *post-hoc* test).

Numerous neuropathological studies have revealed the presence of lower interneuron density in postmortem tissues from individuals with ASDs and hypoactive parvalbumin (PV)^+^ interneurons in various ASD animal models (19, 20). Based on these findings, we examined the density of PV^+^ interneurons and neuronal activities across hippocampal subfields in our animal models (**Supplemental Fig. 8**). However, we did not find any changes in interneuron density or neuronal activity (assessed via c-FOS immunostaining) in adult *Mdga1*^Y636C/E751Q^ KI or *Mdga1*-cKO mice. Given the previous report that MDGA1 is robustly expressed in upper cortical layer neurons (21), we examined whether *Mdga1*^Y636C/E751Q^ KI mice exhibited altered targeting of cortical layer II/III neurons, as seen in *Mdga1*-cKO mice (see accompanying paper). Immunohistochemical analyses performed using the cortical layer II/III-specific markers, Wfs1, calbindin (CB), and Brn2, revealed no change in the density of CB^+^ or Brn2^+^ neurons in somatosensory cortex layer II/III of adult male *Mdga1*^Y636C/E751Q^ KI mice (**Supplemental Fig. 9**), which contrasted with the results obtained in *Mdga1*-cKO mice (see accompanying paper). Our data collectively demonstrate that the Y636C/E751Q substitution induces a sexually dimorphic loss-of-function for MDGA1 that is reflected in both electrophysiological and behavioral phenotypes; the situation is similar to that seen in *Mdga1*-cKO neurons, but does not depend on the developmental trajectory of interneurons.

### Synapsin II is hypophosphorylated in adult male *Mdga1*-cKO and *Mdga1*^Y636C/E751Q^ KI mice

To explore the mechanisms through which MDGA1 loss-of-function induces ASD-like phenotypes, we attempted proteomic and phosphoproteomic analyses of hippocampal tissues from embryonic mice (E18), adult (P60) male *Mdga1*-cKO mice, male *Mdga1*^Y636C/E751Q^ KI mice, and littermate control embryos or mice. The proteomic analysis showed that the MDGA1 protein level was significantly reduced in *Mdga1*-cKO mice but not in *Mdga1*^Y636C/E751Q^ KI mice (**Supplemental Fig. 10; Supplemental Tables 1** and **2**). Phosphopeptides were enriched in tryptic peptides using immobilized metal ion affinity chromatography (IMAC) and subjected to LC-MS/MS (**Fig. 5A**; **Supplemental Tables 3** and **4**). Our analysis of adult male *Mdga1*-cKO mice versus control mice identified 28 differentially expressed phosphopeptides (DEPPs, *p* < 0.05; **Supplemental Table 4**) that exhibited significant changes (Log_2_FC > 0.4 or Log_2_FC < -0.4), and further revealed that they belong to 27 differentially expressed proteins (DEPs). In contrast, our analysis of adult male *Mdga1*^Y636C/E751Q^ KI mice identified 83 DEPPs belonging to 75 DEPs (**Fig. 5B** and **5C**; **Supplemental Table 5**). Gene Ontology (GO) analyses performed using Metascape showed that the DEPs of both *Mdga1*-cKO and *Mdga1*^Y636C/E751Q^ KI mice were enriched for GO terms connected with synapse-related functions (**Fig. 5D**). A substantial fraction of the DEPs from adult male *Mdga1*-cKO mice and a smaller proportion from adult male *Mdga1*^Y636C/E751Q^ KI mice (33.3% and 3%, respectively) belonged to the SynGO protein category (**Fig. 5E–5G**). Among the DEPs, synapsin II (SynII) and OUT domain-containing protein 5 (Otud5) were commonly downregulated in adult male *Mdga1*-cKO and *Mdga1*^Y636C/E751Q^ KI mice (**Fig. 5H** and **5I**).

We next used the DEPs to perform a STRING analysis of protein-protein interactions (PPIs) (**Fig. 5J** and **5K**). Intriguingly, the phosphorylation levels of SynII were clearly decreased in adult male *Mdga1*-cKO and adult male *Mdga1*^Y636C/E751Q^ KI mice. Synaptic vesicle clustering and maintenance of synapse structure were identified as the most relevant biological processes in mice of both genotypes; the relevant DEPs included SynI, SynII, Piccolo, Bassoon, and RIMS1 (**Fig. 5K**). Notably, the phosphorylation levels of these proteins were not significantly altered in our phosphoproteomic analysis of E18 embryos from *Mdga1*^Y636C/E751Q^ KI mice (**Supplemental Fig. 11** and **12**; **Supplemental Table 6**). Together, the data from our extensive proteomics analyses revealed that there are marked developmental stage-related differences in the DEPP profiles, DEP profiles, and phosphorylation levels of some presynaptic proteins, including Syn proteins, that might form the basis for the observed sexually dimorphic abnormalities in MDGA1 functions (see below).

### Bazedoxifene rescues reduced the SynII phosphorylation, impaired communicative behavior, and altered GABAergic synaptic transmission in adult male *Mdga1*^Y636C/E751Q^ KI mice

Given that major phenotypes were observed exclusively in male *Mdga1*-cKO and *Mdga1*^Y636C/E751Q^ KI mice, and males and females exhibit differential sensitivity to the organizational effects of gonadal hormones during the pre-pubertal period (22), we hypothesized that altering estradiol levels in male *Mdga1*^Y636C/E751Q^ KI would normalize their electrophysiological and behavioral abnormalities. We utilized bazedoxifene (BZD), which is a third-generation selective estrogen receptor modulator that is brain blood barrier (BBB)-penetrant and Food and Drug Administration (FDA)-approved for treating menopausal symptoms (23-25). We acutely injected adult male *Mdga1*^Y636C/E751Q^ KI mice with a single dose of BZD (4.1 mg/kg) and, 6 hours later, performed semi-quantitative immunoprecipitation analyses using phosphoserine antibodies (recognizing phosphorylated Syn residues) to measure changes in Syn protein phosphorylation, and applied a subset of behavioral tests to measure ASRs, PPIs and USVs (**Fig. 6A**). As hypothesized, administration of BZD rescued the reduced protein phosphorylation in isoforms of SynII (but not SynI), the downregulation of GABAergic synaptic proteins (Nlgn2, GABA_A_Rγ2, and VGAT; **Fig. 6B–6E**), and several behavioral abnormalities, all of which returned to the levels seen in littermate control or female counterpart mice (**Fig. 6F–6K**). More specifically, a subset of altered USV syllable types (including complex trill) and ASRs, but not PPIs, were rescued by BZD treatment of adult male *Mdga1*^Y636C/E751Q^ KI mice (**Fig. 6H–6K**). In addition, the enhanced GABAergic synaptic strength in the distal dendrites of the CA1 pyramidal neurons from adult male *Mdga1*^Y636C/E751Q^ KI mice was restored to control levels upon BZD treatment (**Fig. 6L–6P**). These results provide the first evidence of a druggable target (i.e., MDGA1) that can be used to alter communication deficits in an animal model of autistic-like features. Viewed together, our results highlight different combinations of synaptic, behavioral and hormonal mechanisms that can produce ASD-like manifestations and suggest that MDGA1 could be a reliable biomarker for studying a putative biological basis for the male preponderance of ASDs.

## Discussion

The current study complements the accompanying study by demonstrating that one of our ASD-associated *MDGA1* variant pairs induces both gain-of-function and loss-of-function phenotypes in adult mice, resulting in autistic-like behavioral abnormalities and imbalanced excitation/inhibition (E/I) ratios in hippocampal CA1 pyramidal neurons. *Mdga1*^Y636C/E751Q^ KI pups emitted fewer isolation-induced USVs, and thus mirrored the phenotypes of *Mdga1*-cKO pups (see accompanying paper); this suggests that the endogenous MDGA1 protein may compensate for the defects induced by the overexpressed mutant protein. The Y636C/E751Q substitution did not alter the hippocampal level of MDGA1, despite our *in silico* prediction that these changes would exert a subtle destabilizing effect (see accompanying paper). Remarkably, adult male *Mdga1*^Y636C/E751Q^ KI mice exhibited increased USV calls with abnormal USV syllable composition; this behavior contrasts with the decreased USV calls observed in adult male *Mdga1*-cKO mice, which is typical of other ASD model mice. The increase of USV calls in adult male *Mdga1*^Y636C/E751Q^ KI mice might reflect a gain-of-function effect that leads to an increase in biologically irrelevant USV calls. Further studies are warranted to unravel the mechanisms by which the potentially marginal destabilization of MDGA1 by the Y635C/E756Q substitution leads to either loss-of-function or gain-of-function effects, and to determine whether the V116M/A688V substitution causes a similar change of MDGA1 protein levels.

Our proteomic analyses indicated that *Mdga1*-cKO and *Mdga1*^Y636C/E751Q^ KI neurons display reduced phosphorylation of presynaptic SynII, which is likely desynchronize neurotransmitter release at GABAergic synapses. This is consistent with the observation that the action of MDGA1 is transsynaptically linked to presynaptic neurons (15). We speculate that MDGA1-mediated negative transsynaptic regulation occurs via precisely tethered APP protein, which is further coupled with the activity of SynII proteins to orchestrate GABA release at the synaptic cleft that is designed to maximize synaptic strength via the precise alignment of pre- and postsynaptic compartments (26, 27). It is strategically advantageous for MDGA1 to adopt a compact triangular conformation that facilitates mechanical stabilization and efficient transsynaptic signaling (26, 28). It would be interesting to explore whether the circuit-specific introduction of extracellularly stabilized MDGA1 protein or drugs that selectively boost specific SynII phosphorylation site(s) could have beneficial effects on ASDs in the context of various neurodevelopmental trajectories.

Our study raises several questions/issues that remain to be addressed. Whereas adult female *Mdga1*-cKO and *Mdga1*^Y636C/E751Q^ KI mice displayed no abnormal behavioral or electrophysiological phenotype, juvenile female *Mdga1*-cKO and *Mdga1*^Y636C/E751Q^ KI mice exhibited electrophysiological phenotypes similar to those of their male counterparts. Although dimorphic phenotypes have been reported in numerous ASD models (29), definitive explanations are lacking. The female protective effect theory of ASDs (30, 31) is compelling, since a sudden peak in gonadal hormone levels during the prepubertal period of female mouse development appears to contribute to an anti-inflammatory state that further regulates hippocampal CA1 GABAergic synapse organization (32, 33). We herein demonstrate that BZD may be useful for treating certain aspects of ASDs, and our findings further imply that estradiol signaling may act on synaptic disinhibition to normalize SynII activities (34).

Moreover, GABAergic synaptic inhibition is essential for controlling the window of the critical period of plasticity (35) and its precocious closure is linked to ASDs (36, 37), giving rise to the speculation that MDGA1 is a key regulatory factor in the development of critical-period plasticity. The effects of BZD administration should be systematically examined in other ASD model animals that exhibit sexually dimorphic abnormalities. We also need to better understand the synaptic and circuit-level actions of BZD and how long its rescue effects last during development. Such studies would help inform the optimization of BZD dosing and administration frequency, with the goal of avoiding potentially adverse side effects (38). Given the significant expression of MDGA1 in astrocytes, it would also be interesting to establish whether both neuronal and astrocytic MDGA1 are engaged in shaping the critical periods that determine sexual dimorphism during development across diverse brain areas (39-41). Another future challenge will be to determine the cell types, circuits, and/or regions that are primarily involved in eliciting various behavioral manifestations of *Mdga1*-cKO and *Mdga1*^Y636C/E751Q^ KI mice. Notably, overexpression or deletion of MDGA1 in mPFC neurons failed to exert significant effects on synaptic properties and mouse behavior, partly in line with the low-level expression of MDGA1 in the mPFC. Considering the contributions of the hippocampus to the social and cognitive deficits in ASDs (42), our results obtained from the hippocampal CA1 GABAergic circuits are compellingly relevant to ASD pathophysiology. To gain a better understanding of ASD biology, the investigation of MDGA1 functions should be extended beyond the hippocampus and mPFC to other regions related to regulating ASD-like behaviors, such as the thalamus (43, 44), with the goals of addressing the canonical/non-canonical roles of MDGA1 and establishing its utility as a genuine biomarker of ASD. Importantly, in addition to homozygous *Mdga1*^Y636C/E751Q^ KI mice, heterozygous *Mdga1*^Y636C/E751Q^ KI mice should be studied to elucidate the detailed abnormalities relevant to ASD patients.

## Methods

### Animals

All mice were maintained and handled in accordance with protocols (DGIST-IACUC-23112809-0003) approved by the Institutional Animal Care and Use Committee of DGIST under standard, temperature-controlled laboratory conditions. Mice were kept on a 12:12-h light/dark cycle (lights on at 7:00 am) and received water and food *ad libitum*. All experimental procedures were performed on male mice, using littermate control without Cre expression. Conditional *Mdga1*-KO mice was previously described (15, 45). Pregnant rats (Daehan Biolink) were used to prepare *in vitro* cultures of dissociated hippocampal neurons. *Mdga1*^Y636C/E751Q^-KI mice were generated by Biocytogen Pharmaceuticals (Beijing) based on CRISPR/Cas9 approach. To generate *Mdga1^Mut/+^* mice, the candidate sgRNAs, located in the intron9 and intron13 of *Mdga1*, were searched by the CRISPR design tool (http://www.sanger.ac.uk/htgt/wge/) and then were screened for on-target activity using a Universal CRISPR Activity Assay (UCA^TM^, Biocytogen Pharmaceuticals (Beijing Co., Ltd). A gene-targeting vector containing a 5’ homologous arm, donor fragment (containing the mutations of p.Tyr636Cys; TAC>TGC and p.Glu751Gln; GAG>CAG), and 3’ homologous arm was used as a template to repair the double-stranded breaks (DSBs) generated by Cas9/sgRNA. The T7 promoter sequence was added to the Cas9 or sgRNA template by PCR amplification *in vitro*. The Cas9 mRNA, targeting vector, and sgRNAs were co-injected into the cytoplasm of one-cell stage fertilized C57BL/6N eggs. The injected zygotes were transferred into oviducts of Kunming pseudo-pregnant females to generate F0 mice. F0 mice with expected genotype confirmed by tail genomic DNA PCR and sequencing were mated with C57BL/6N mice to establish germline-transmitted F1 heterozygous mice. F1 heterozygous mice were genotyped by tail genomic PCR, southern blot, and DNA sequencing.

### Antibodies

The following commercially available antibodies were used: mouse monoclonal anti- APZP (clone 22C11; Millipore; Cat# MAB348; RRID: AB_94882); mouse monoclonal anti-β-actin (clone AC-74; Sigma-Aldrich; Cat# sc-47778; RRID: AB_476743); mouse monoclonal anti-Nlgn1 (Synaptic Systems; Cat# 129 111; RRID: AB_887747); rabbit polyclonal anti-Nlgn2 (Synaptic Systems; Cat# 129 202; RRID: AB_993011); guinea pig polyclonal anti-VGLUT1 (Millipore; Cat# ab5905; RRID: AB_2301751); mouse monoclonal anti-gephyrin (Synaptic Systems; Cat# 147 011; RRID: AB_887719); rabbit polyclonal anti-VGAT (Synaptic Systems; Cat# 131 003; RRID: AB_887869); rabbit polyclonal anti-GABA_A_ receptor γ2 (Synaptic Systems; Cat# 224 003; RRID: AB_2263066); mouse monoclonal anti-GAD67 (clone 1G10.2; Millipore; Cat# MAB5406; RRID: AB_2278725); mouse monoclonal anti- NMDAR1 (Clone 54.1; Millipore; Cat# MAB363; RRID: AB_94946); rabbit polyclonal anti-PSD-95 (goat polyclonal anti-EGFP (Rockland; Cat# 600-101-215; RRID: AB_218182); chicken polyclonal anti-EGFP (Aves Labs; Cat# GFP-1020; RRID: AB_10000240); mouse monoclonal anti-HA (clone 16B12; BioLegend; Cat# 901501; RRID: AB_2565006); rabbit monoclonal anti-HA (clone C29F4; Cell Signaling Technology; Cat# 3724; RRID: AB_1549585); rat monoclonal anti-somatostatin (clone YC7; Millipore; Cat# MAB354; RRID: AB_2255365); mouse monoclonal anti-PV (clone PV235; Swant; Cat# 235; RRID: AB_10000343); rabbit monoclonal anti-c-Fos (clone 9F6; Cell Signaling Technology; Cat# 14609; RRID: AB_2798537); rabbit polyclonal anti-phosphoserine (Millipore; Cat# AB1603; RRID: AB_390205); mouse monoclonal anti-Synapsin I (Synaptic Systems; Cat# 106 011; RRID: AB_2619722); mouse monoclonal anti-Synapsin II (Synaptic Systems; Cat# 106 211; RRID: AB_2619774); rabbit polyclonal anti-phospho-Synapsin II (Ser425) (Thermo Fisher Scientific; Cat# PA5-64855; RRID: AB_2663626); mouse monoclonal anti-Calbindin (Swant; Cat# 300; RRID: AB_10000347); rabbit polyclonal anti-Wfs1 (Proteintech; Cat# 11558-1-AP; RRID: AB_2216046); chicken polyclonal anti-Tbr1 (Millipore; Cat# AB2261; RRID: AB_10615497); rabbit polyclonal anti-Brn2 (Abcam; Cat# ab94977; RRID: AB_10615497); and Cy3-donkey anti-human IgG antibodies (Jackson ImmunoResearch; Cat# 709-165- 149; RRID: AB_2340535). The following antibody was previously described: rabbit polyclonal antipan-SHANK (1172; RRID: AB_2810261) and rabbit polyclonal anti-PSD-95 (JK016; RRID: AB_2722693) (46).

### Immunohistochemistry, confocal microscopy imaging, and analyses

P60 mice were anaesthetized and immediately perfused, first with PBS for 3 min, and then with 4% paraformaldehyde for 5 min. Brains were dissected out, fixed in 4% paraformaldehyde overnight, and sliced into 30-um-thick coronal sections using a vibratome (Model VT1200S; Leica Biosystems).

Sections were permeabilized by incubating with 2% Triton X-100 in PBS containing 5% bovine serum albumen and 5% horse serum for 30 min. For immunostaining, sections were incubated for 8–12 h at 4°C with primary antibodies diluted in 0.1% Triton X-100 in PBS containing bovine and horse serum. The following primary antibodies were used: anti-VGLUT1 (1:300), anti-PSD-95 (1:300), anti-VGAT (1:300), anti-gephyrin (1:500), anti-PV (1:300), anti-SST (1:50), anti-c-Fos (1:400), anti-Brn2 (1:100), anti- Wfs1 (1:100), and anti-Calb (1:100). Sections were washed three times in PBS and incubated with appropriate Cy3- or FITC-conjugated secondary antibodies (Jackson ImmunoResearch) for 2 h at room temperature. After three washes with PBS, sections were mounted onto glass slides (Superfrost Plus; Fisher Scientific) with VECTASHIELD mounting medium (H-1200; Vector Laboratories). Images were acquired by confocal microscopy (LSM800; Zeiss). Synaptic puncta were quantified using MetaMorph software, and their density and average area were measured. For immunohistochemical analysis of electroporated mice (P21), the mice were heart-perfused with 4% paraformaldehyde.

Brains were collected, embedded in 3% low melting-temperature agarose, and sectioned at 100 μm thickness with a vibratome (VT1200S; Leica). The sections were incubated with primary antibodies (anti-EGFP, 1:2000; anti-HA, 1:1000) in PBS containing 0.1% Triton X-100, 0.1% BSA, and 2.5% goat serum, and then with fluorescent secondary antibodies (Alexa 488 and Alexa 647). Slices were mounted in Antifade mounting medium with DAPI (VECTASHIELD) and images were acquired by confocal microscopy (A1R; Nikon).

### Preparation of adeno-associated viruses and titration

HEK293T cells were co-transfected with the indicated AAV vectors and pHelper and pAAV1.0 (serotype 2/9) vectors. 72 hours later, transfected HEK293T cells were collected, lysed, and mixed with 40% polyethylene glycol and 2.5 M NaCl, and centrifuged at 2,000 × g for 30 min. The cell pellets were resuspended in HEPES buffer (20 mM HEPES; 115 mM NaCl, 1.2 mM CaCl_2_, 1.2 mM MgCl_2_, 2.4 mM KH_2_PO_4_) and an equal volume of chloroform was added. The mixture was centrifuged at 400 × g for 5 min, and concentrated three times with a Centriprep centrifugal filter (Millipore) at 1,220 × g for 5 min each and with an Amicon Ultra centrifugal filter (Millipore) at 16,000 × g for 10 min. Before the titration of AAVs, contaminating plasmid DNA was eliminated by treating 1 μl of concentrated, sterile-filtered AAVs with 1 μl of DNase I (Sigma-Aldrich) for 30 min at 37°C. After treatment with 1 μl of stop solution (50 mM ethylenediaminetetraacetic acid) for 10 min at 65°C, 10 μg of protease K (Sigma-Aldrich) was added and AAVs were incubated for 1 h at 50°C. Reactions were inactivated by incubating samples for 20 min at 95°C. The final virus titer was quantified by qRT-PCR detection of EGFP sequences and subsequent reference to a standard curve generated using the pAAV-T2A-EGFP plasmid. All plasmids were purified using a Plasmid Maxi Kit (Qiagen GmbH).

### Stereotactic surgery and virus injections

Adult (P30) mice were anesthetized by intraperitoneal injection of a saline-based 2% Avertin solution (2,2,2-tribromoethyl alcohol dissolved in tert- amylalcohol [Sigma]), and their heads were fixed in a stereotactic apparatus. Recombinant AAV virus was injected into the hippocampal CA1 region (coordinates: AP -2.1 mm, ML ± 1.3 mm, and DV 1.8 mm) with a Hamilton syringe at a flow rate of 100 nl/min (injected volume, 300 nl) using a Nanoliter 2010 Injector (World Precision Instruments). Each injected mouse was restored to its home cage for 2- 4 weeks and used subsequently for immunohistochemical analyses, electrophysiological recordings, or behavioral analyses.

### Electrophysiology

Hippocampal slices (300 μm) were prepared from mice aged 10–12 days, 18–20 days, or 8–10 weeks. Following anesthesia with isoflurane, the mice were euthanized, and their brains were swiftly extracted and placed in a chilled solution with low calcium and high magnesium levels, which was oxygenated (95% O_2_ and 5% CO_2_). This solution consisted of 3.3 mM KCl, 1.3 mM NaH_2_PO_4_, 26 mM NaHCO_3_, 11 mM D-glucose, 0.5 mM CaCl_2_, 10 mM MgCl_2_, and 211 mM sucrose.

Hippocampal slices were prepared using a vibratome (VT1000S; Leica) and moved to a storage chamber filled with oxygenated artificial cerebrospinal fluid (aCSF) containing 124 mM NaCl, 3.3 mM KCl, 1.3 mM NaH_2_PO_4_, 26 mM NaHCO_3_, 11 mM D-glucose, 2 mM CaCl_2_, and 1 mM MgCl_2_. Slices were incubated at 30°C for at least 60 minutes prior to experimentation. Subsequently, slices were moved into the recording chamber and subjected to constant perfusion with standard aCSF that was oxygenated with a mixture of 95% O_2_ and 5% CO_2_. All experiments were performed at 32°C, and slices were used within 4 h. Only cells with access resistance (Ra) smaller than 30 MΩ were analyzed. All the recordings were conducted using a Multiclamp 700B amplifier and DigiData 1550B Digitizer. To assess miniature inhibitory postsynaptic currents (mIPSCs), whole cell recordings were performed from hippocampal CA1 pyramidal neurons using glass pipettes (3–5 MΩ) filled with a solution containing 145 mM CsCl, 5 mM NaCl, 10 mM HEPES, 10 mM EGTA, 4 mM Mg-ATP, and 0.3 mM Na- GTP, adjusted to pH 7.2–7.3 with CsOH. Cellular voltage was clamped at -70 mV. mIPSCs were isolated by inhibiting NMDAR, AMPAR, and Na^+^ channel using external application of 50 μM D-AP5, 10 μM CNQX, and 1 μM tetrodotoxin. For measurement of evoked inhibitory postsynaptic currents (eIPSCs) and delayed asynchronous IPSCs (aIPSCs), patch pipettes were filled with an internal solution consisting of 130 mM Cs-methanesulfonate, 5 mM TEA-Cl, 8 mM NaCl, 0.5 mM EGTA, 10 mM HEPES, 4 mM Mg-ATP, 0.4 mM Na-GTP, 1 mM QX-314, and 10 mM disodium phosphocreatine, adjusted to pH 7.2–7.3 with CsOH. Cells were voltage-clamped at 0 mV. Electrical stimulation was applied using a concentric bipolar electrode (FHC), placed on the *stratum radiatum* (SR) or *stratum lacunosum moleculare* (SLM) in the hippocampal CA1 to record somatic inhibition and dendritic inhibition respectively. eIPSCs and aIPSCs were isolated by inhibiting NMDAR, and AMPAR using external application of 50 μM D-AP5, and 10 μM CNQX. To record eIPSCs input-output (eIPSCs I-O), electrical stimulation was applied ranged 10 to 60 μA with 10 μA increments. Average eIPSCs I-O were measured from three consecutive sweeps. For assessing paired-pulse ratios (PPRs), pairs of stimuli were administered at 10 Hz frequency. PPRs were calculated as 2^nd^ eIPSC/1^st^ eIPSC. Average PPRs were analyzed from three consecutive sweeps. To assess delayed aIPSCs, 40 Hz train stimulation was applied. Synaptic events during 1 second area after train stimulation were quantified, normalized to the 1 second area before the train stimulation.

### Mouse behavioral tests

Male and female mice aged 7–9 weeks were used for all behavioral tests. Tests were conducted in a soundproofed room under dim lighting (< 5 lux). All behavioral examinations were conducted in a controlled environment and mice were acclimated to the experimental room prior to testing. The order of testing was randomized, and experiments conducted at consistent timeframes to minimize potential circadian influences.

*<u>1. Open-field test</u>.* Mice were placed in a white acrylic open-field box (40 × 40 × 40cm) and allowed to freely explore the environment for 10 min in dim light (< 5 lux). The distance traveled and time spent in the center zone by freely moving mice were recorded with a top-view infrared camera and analyzed using EthoVision XT 10.5 software (Noldus).
*<u>2. Y-maze test.</u>* A Y-shaped white acrylic maze with three 35 cm long arms at a 120ʹ angle from each other was used. Mice were introduced into the maze and allowed to explore freely for 8 min. A move was counted as an entry when all four limbs of the mouse were within the arm. Movement of mice was recorded with a top-view infrared camera and analyzed using EthoVision XT 10.5 software.
*<u>3. Novel object-recognition test.</u>* An open-field chamber was used for this experiment. Mice were habituated to the chamber for 10 min. For training sessions, two identical objects were placed in the center of the chamber at regular intervals and mice allowed to explore the objects for 10 min. After the training session, mice were returned to their home cages for 24 h. For novel object recognition tests, one of the two objects was replaced with a new object, which was placed in the same position in the chamber. Mice were returned to the chamber and allowed to explore freely for 10 min. Movement was recorded using an infrared camera and the number and duration of contacts analyzed using EthoVision XT 10.5. The discrimination index represents the difference between time spent exploring novel and familiar objects during the test phase.
*<u>4. Elevated plus-maze test.</u>* The white acrylic maze with two open arms (30 × 5 × 0.5 cm) and two closed arms (30 × 5 × 30 cm) was elevated by 75 cm over the floor. Mice were individually placed at the center of the elevated plus-maze and allowed to freely move for 5 min. All behaviors were recorded with a top-view infrared camera, and the time spent in each arm and number of arm entries analyzed using EthoVision XT 10.5.
*<u>5. Light and dark box transition test.</u>* The apparatus consisted of a roofless box divided into a closed dark chamber and a brightly illuminated chamber. A small entrance allowed free travel between the two chambers. The light chamber was illuminated at 350 lux. Movement of mice was recorded using a top-view infrared camera and EthoVision XT 10.5 applied for analysis of the time spent in each chamber and the number of transitions.
*<u>6. Three-chamber test.</u>*The testing apparatus consisted of a white acrylic box divided into three chambers (20 × 40 × 22 cm each) with small openings on the dividing walls. Wire cups were employed to enclose social conspecifics in the corners of both side chambers. Mice were placed in the central chamber for a 10 min habituation period. Following this time, an age-matched social conspecific was placed in the wire cup on the left side-chamber and sociability of the subjects assessed by measuring subject exploration times for the enclosed conspecific and empty cup during the second 10 min session. An exploration was counted when the subjects placed their nose in the vicinity of the wire cups. In the final 10 min session, a new social conspecific was placed into the empty wire cup on the right side-chamber and social recognition assessed by measuring subject exploration times of the familiar and novel conspecifics in the wire cup by the subject.
*<u>7. Prepulse inhibition test.</u>* Mice were individually acclimated to the experimental room for a minimum of 30 min under standard housing conditions with reduced extraneous noise. Each mouse was placed in a sound-attenuating chamber equipped with startle response measurement apparatus (SR-LAB- Startle Response System; SD Ins.), ensuring minimal movement during testing. On day 1, after a 5- min habituation period in the absence of stimuli, mice were exposed to a series of startle-eliciting stimuli (from 0 to 120 dB) to familiarize them with the startle response. On day 2, for the PPI trials, prepulse stimuli (73, 76, 79, or 82 dB) were presented before the startle-eliciting stimuli (120 dB). The order and inter-trial intervals of prepulse and startle stimuli were randomized, including control trials with no prepulse. Startle responses were recorded for each trial using PPI test software. Data analysis involved calculating the percentage PPI for each prepulse intensity using the formula: *PPI* (%) = (*R_pulse_*−*R_pre_*−*pulse*)/*R_pulse_* × 100 (*R_pulse_* is the average response magnitude in the absence of a prepulse and *R_pre_*−*pulse* the average response magnitude in the presence of a prepulse).
*<u>8. Ultrasonic vocalization test</u>*. Following a 3-day period of single housing, each adult mouse was individually placed in a sound-attenuating chamber equipped with a microphone for USV recording. To introduce a stimulus for vocalization, a conspecific of the opposite sex, rendered anesthetized, was gently introduced into the chamber. The interactions and vocalizations of the focal mouse were recorded for a duration of 10 min. In experiments involving pups, sessions were conducted on postnatal days P3, P6, P9, and P12. Each pup was temporarily separated from the dam and placed in a cage with clean bedding on a heat pad maintained at 37 ^0^C to ensure body temperature regulation and its emitted ultrasonic vocalizations meticulously measured for 10 min. USV recordings were subjected to detailed analysis using DeepSqueak software (reference) for the detection and categorization of ultrasonic vocalizations. This software facilitated the extraction of key parameters, such as call frequency, duration, and type. Calls were systematically categorized based on their spectrotemporal features.
*<u>9. Forced swim test.</u>* Mice were individually placed in a glass cylinder (15ⅹ30 cm) containing water (24 ± 1 °C; depth, 15 cm). All mice were forced to swim for 6 min, and the duration of immobility measured during the final 4 min of the test. The latency to immobility from the start of the test (delay between the start of the test and appearance of the first bout of immobility, defined as a period of at least 1 s without any active escape behavior) and duration of immobility (defined as the time not spent actively exploring the cylinder or trying to escape from it) were measured. Immobility time was defined as the time the mouse spent floating in the water without struggling, making only minor movements that were strictly necessary to maintain its head above water.
*<u>10. Tail-suspension test.</u>*Mice were individually subjected to the tail suspension test, involving suspension by adhesive tape affixed approximately 1-2 cm from the tail tip. The testing apparatus consisted of a standardized setup and mice were allowed to hang freely. The total test duration was set at 5 min during which mice were examined for despair-related behaviors. The immobility time was recorded for the final 4 min of the test. Immobility was operationally defined as the absence of limb or body movement, and the recorded duration represented the time spent suspended without exhibiting active escape behaviors or exploratory movements. Additionally, latency to immobility was measured as the period during which the mouse did not engage in any active escape behavior for at least 1 second.
*<u>11. Marble burying test.</u>* Mice were individually acclimated to the experimental room for at least 30 min prior to testing. Standard mouse bedding was placed in home cages to provide a familiar environment. For the marble burying test, a 5 cm layer of fresh bedding was uniformly spread in the Plexiglas cage. Next, 15 glass marbles were evenly distributed on the bedding surface. Each mouse was introduced into the cage and allowed free exploration for a duration of 30 min. During the marble burying phase, the number of marbles buried (defined as being at least two-thirds covered by bedding) was counted and quantified.

### Proteomics

*<u>1. Peptide preparation and phosphopeptide enrichment</u>.* Hippocampi from male control, *Mdga1*-cKO, or *Mdga1*^Y636C/E751Q^ KI mice (4 replicates) of adult (P60) and E18 mouse embryos were lysed with 1X SDS buffer (5% SDS and 50 mM triethylammonium bicarbonate, pH 8.5). The lysates were digested using an S-trap method with trypsin, according to the provided protocol. The digested peptides were labeled with 18-plex TMT isotopes (Thermo Fisher Scientific), dried by Speed-Vac, and desalted with Pierce peptide desalting spin columns (Thermo Fisher Scientific). The combined samples were fractionated into 20 fractions by basic reverse phase liquid chromatography. Of the sample, 5% was used for total proteome analysis and then the remaining 95% was reserved for phosphoproteomic analysis. The remaining latter portion was combined into 10 fractions for phosphopeptide enrichment. First, Ni-NTA magnetic agarose beads were washed three times with DW and then incubated with 100 mM EDTA (pH 8.0) for 30 minutes on a rotator. Next, 100 mM FeCl_3_ solution was added and the mixture was incubated with rotation for 30 minutes. The prepared Fe^3+^-NTA beads were washed with DW and incubated overnight with each sample in 80% ACN with 0.1% TFA, at 4°C on a rotator. The bead-bound phosphopeptides were eluted with elution buffer (50% ACN in 1% ammonium hydroxide), promptly acidified to pH 3.5–4.0 using 10% TFA, and vacuum dried.
*<u>2. LC-MS/MS and data analysis.</u>* LC-MS/MS analysis was performed using an UltiMate 3000 RSLCnano system (Thermo Fisher Scientific) coupled to a Orbitrap Fusion Lumos mass spectrometer (Thermo Fisher Scientific). Mobile phases A and B were composed of 0 and 95.0% acetonitrile containing 0.1% formic acid, respectively. The LC gradient was applied at a flow rate of 250 nL/min during 120 min for peptide separation. The Orbitrap Fusion Lumos was operated in data-dependent mode, and the MS2 scans were performed with HCD fragmentation (37.5% collision energy). MS/MS spectra were identified and quantified using the Integrated Proteomics Pipeline software with the UniProt mouse database and the following search parameters: precursor mass tolerance, 20 ppm; fragment ion mass tolerance, 200 ppm; two or more peptide assignments for protein identification at a false positive rate < 0.01; and TMT reporter ion mass tolerance, 20 ppm. For phosphopeptide identification, phosphorylation of serine, threonine, and tyrosine was set as the differential modifications, with a maximum of three additional modifications permitted. Statistical analysis was conducted using the Perseus software (version 1.6.15). The expression levels of proteins and phosphopeptides between samples were compared with Welchʹs t-test, with the *p* value set at < 0.05. The raw MS data files for total proteomics have been deposited to the MassIVE repository with identifier PXD057134 (https://proteomescentral.proteomaxchange.org/cgi/GetDataset?ID=PXD057134) for data from *Mdga1*- cKO mice and identifier PXD057136 (https://proteomescentral.proteomexchange.org/cgi/GetDataset?ID=PXD057136) for data from *Mdga1*^Y636C/E751Q^ KI mice.

### Statistical analysis

No statistical method was used to pre-determine sample sizes. Rather, the sample sizes were selected based on previous studies published in the field (see Life Science Reporting Summary for references). Animals in the same litter were randomly assigned to the different treatment groups in the various experiments. All statistical analyses were performed using GraphPad Prism 7 software (RRID: SCR_002798). The normality of distributed data was determined using the Shapiro-Wilk normality test. Normally distributed data were compared using Student’s t- test or the analysis of variance (ANOVA) test, and non-normally distributed data by the Mann– Whitney *U* test, non-parametric ANOVA with Kruskal–Wallis test followed by *post hoc* Dunn’s multiple comparison test, or non-parametric ANOVA with *post hoc* Tukey’s multiple comparison test. If a single value made the data distribution non-normal and was found to be a significant outlier (*p* < 0.05) by Grubb’s test, it was regarded as an outlier. **Supplementary Table 11** presents detailed statistics.

## Author contributions

Conceptualization: Jaewon K.; Data curation: S.K., Jinhu K., B.K., H.K., Y.Y. and H.J.L.; Funding acquisition: S.K., H.K., J.Y.K., J.W.U. and Jaewon K.; Investigation: S.K., Jinhu K., B.K., H.K.,Y.Y. and

H.J.L.; Methodology: S.K., Jinhu K., B.K., H.K., Y.Y., H.J.L. and J.L.; Supervision: J.Y.K., J.W.U. and Jaewon K.; Writing – original draft: Jaewon K.; Writing – review & editing: Jaewon K.

## Acknowledgments

We thank Jinha Kim (DGIST) for technical assistance. This study was supported by National Creative Research Initiative Program of the Ministry of Science and ICT (RS-2022-NR070708 to Jaewon K.), the National Research Foundation of Korea (NRF) funded by the Korea Government (2022R1C1C200003499 to S.K.; RS-2024-00339642 to H.K.; RS-2024-00399031 to J.Y.K.; and 2023R1A2C2002535 to J.W.U.), and the Korea Basic Science Institute (A423200 to J.Y.K.).

## Conflict-of-interest statement

The authors have declared that no conflict of interest exists.

## Notes

### Competing Interest Statement

The authors have declared no competing interest.

## References

1. Südhof TC. Towards an Understanding of Synapse Formation. Neuron. 2018;100(2):276-93.

2. Kim HY, Um JW, and Ko J. Proper synaptic adhesion signaling in the control of neural circuit architecture and brain function. Prog Neurobiol. 2021;200:101983.

3. Südhof TC. The cell biology of synapse formation. J Cell Biol. 2021;220(7):e202103052.

4. Kim J, Park D, Seo NY, Yoon TH, Kim GH, Lee SH, et al. LRRTM3 regulates activity-dependent synchronization of synapse properties in topographically connected hippocampal neural circuits. Proc Natl Acad Sci U S A. 2022;119(3):e2110196119.

5. Berns DS, DeNardo LA, Pederick DT, and Luo L. Teneurin-3 controls topographic circuit assembly in the hippocampus. Nature. 2018;554(7692):328–33.

6. Sando R, Jiang X, and Südhof TC. Latrophilin GPCRs direct synapse specificity by coincident binding of FLRTs and teneurins. Science. 2019;363(6429):eaav7969.

7. Kim J, Wulschner LEG, Oh WC, and Ko J. Trans-synaptic mechanisms orchestrated by mammalian synaptic cell adhesion molecules. Bioessays. 2022:e2200134.

8. Um JW, and Ko J. Neural Glycosylphosphatidylinositol-Anchored Proteins in Synaptic Specification. Trends Cell Biol. 2017;27(12):931-45.

9. Connor SA, Elegheert J, Xie Y, and Craig AM. Pumping the brakes: suppression of synapse development by MDGA-neuroligin interactions. Curr Opin Neurobiol. 2019;57:71-80.

10. Kim S, Jang G, Kim H, Lim D, Han KA, Um JW, et al. MDGAs perform activity-dependent synapse type-specific suppression via distinct extracellular mechanisms. Proc Natl Acad Sci U S A. 2024;121(26):e2322978121.

11. Kim JA, Kim D, Won SY, Han KA, Park D, Cho E, et al. Structural Insights into Modulation of Neurexin-Neuroligin Trans-synaptic Adhesion by MDGA1/Neuroligin-2 Complex. Neuron. 2017;94(6):1121-31.

12. Gangwar SP, Zhong X, Seshadrinathan S, Chen H, Machius M, and Rudenko G. Molecular Mechanism of MDGA1: Regulation of Neuroligin 2:Neurexin Trans-synaptic Bridges. Neuron. 2017;94(6):1132-41 e4.

13. Elegheert J, Cvetkovska V, Clayton AJ, Heroven C, Vennekens KM, Smukowski SN, et al. Structural Mechanism for Modulation of Synaptic Neuroligin-Neurexin Signaling by MDGA Proteins. Neuron. 2017;95(4):896-913.

14. Lee K, Kim Y, Lee SJ, Qiang Y, Lee D, Lee HW, et al. MDGAs interact selectively with neuroligin-2 but not other neuroligins to regulate inhibitory synapse development. Proc Natl Acad Sci U S A. 2013;110(1):336-41.

15. Kim J, Kim S, Kim H, Hwang IW, Bae S, Karki S, et al. MDGA1 negatively regulates amyloid precursor protein-mediated synapse inhibition in the hippocampus. Proc Natl Acad Sci U S A. 2022;119(4).

16. Li J, Liu J, Feng G, Li T, Zhao Q, Li Y, et al. The MDGA1 gene confers risk to schizophrenia and bipolar disorder. Schizophr Res. 2011;125(2-3):194-200.

17. Kahler AK, Djurovic S, Kulle B, Jonsson EG, Agartz I, Hall H, et al. Association analysis of schizophrenia on 18 genes involved in neuronal migration: MDGA1 as a new susceptibility gene. Am J Med Genet B Neuropsychiatr Genet. 2008;147B(7):1089-100.

18. Connor SA, Ammendrup-Johnsen I, Kishimoto Y, Karimi Tari P, Cvetkovska V, Harada T, et al. Loss of Synapse Repressor MDGA1 Enhances Perisomatic Inhibition, Confers Resistance to Network Excitation, and Impairs Cognitive Function. Cell Rep. 2017;21(13):3637-45.

19. Contractor A, Ethell IM, and Portera-Cailliau C. Cortical interneurons in autism. Nat Neurosci. 2021;24(12):1648-59.

20. Filice F, Janickova L, Henzi T, Bilella A, and Schwaller B. The Parvalbumin Hypothesis of Autism Spectrum Disorder. Front Cell Neurosci. 2020;14:577525.

21. Takeuchi A, and OʹLeary DD. Radial migration of superficial layer cortical neurons controlled by novel Ig cell adhesion molecule MDGA1. J Neurosci. 2006;26(17):4460-4.

22. Ferri SL, Abel T, and Brodkin ES. Sex Differences in Autism Spectrum Disorder: a Review. Curr Psychiatry Rep. 2018;20(2):9.

23. Zafar E, Maqbool MF, Iqbal A, Maryam A, Shakir HA, Irfan M, et al. A comprehensive review on anticancer mechanism of bazedoxifene. Biotechnol Appl Biochem. 2022;69(2):767-82.

24. Peng L, Luo Q, and Lu H. Efficacy and safety of bazedoxifene in postmenopausal women with osteoporosis: A systematic review and meta-analysis. Medicine (Baltimore). 2017;96(49):e8659.

25. Hill RA, Kouremenos K, Tull D, Maggi A, Schroeder A, Gibbons A, et al. Bazedoxifene - a promising brain active SERM that crosses the blood brain barrier and enhances spatial memory. Psychoneuroendocrinology. 2020;121:104830.

26. Savtchenko LP, and Rusakov DA. The optimal height of the synaptic cleft. Proc Natl Acad Sci U S A. 2007;104(6):1823-8.

27. Biederer T, Kaeser PS, and Blanpied TA. Transcellular Nanoalignment of Synaptic Function. Neuron. 2017;96(3):680-96.

28. Zuber B, Nikonenko I, Klauser P, Muller D, and Dubochet J. The mammalian central nervous synaptic cleft contains a high density of periodically organized complexes. Proc Natl Acad Sci U S A. 2005;102(52):19192-7.

29. Jeon SJ, Gonzales EL, Mabunga DFN, Valencia ST, Kim DG, Kim Y, et al. Sex-specific Behavioral Features of Rodent Models of Autism Spectrum Disorder. Exp Neurobiol. 2018;27(5):321-43.

30. Tsai L, Stewart MA, and August G. Implication of sex differences in the familial transmission of infantile autism. J Autism Dev Disord. 1981;11(2):165–73.

31. Dougherty JD, Marrus N, Maloney SE, Yip B, Sandin S, Turner TN, et al. Can the ʺfemale protective effectʺ liability threshold model explain sex differences in autism spectrum disorder? Neuron. 2022;110(20):3243-62.

32. Mukherjee J, Cardarelli RA, Cantaut-Belarif Y, Deeb TZ, Srivastava DP, Tyagarajan SK, et al. Estradiol modulates the efficacy of synaptic inhibition by decreasing the dwell time of GABA(A) receptors at inhibitory synapses. Proc Natl Acad Sci U S A. 2017;114(44):11763-8.

33. Tabatadze N, Huang G, May RM, Jain A, and Woolley CS. Sex Differences in Molecular Signaling at Inhibitory Synapses in the Hippocampus. J Neurosci. 2015;35(32):11252-65.

34. Ohtani-Kaneko R, Iwafuchi M, Iwakura T, Muraoka D, Yokosuka M, Shiga T, et al. Effects of estrogen on synapsin I distribution in developing hypothalamic neurons. Neurosci Res. 2010;66(2):180-8.

35. Andrade-Talavera Y, Perez-Rodriguez M, Prius-Mengual J, and Rodriguez-Moreno A. Neuronal and astrocyte determinants of critical periods of plasticity. Trends Neurosci. 2023;46(7):566-80.

36. Berger JM, Rohn TT, and Oxford JT. Autism as the Early Closure of a Neuroplastic Critical Period Normally Seen in Adolescence. Biol Syst Open Access. 2013;1.

37. LeBlanc JJ, and Fagiolini M. Autism: a ʺcritical periodʺ disorder? Neural Plast. 2011;2011:921680.

38. Raina PM, Patel P, and Parmar M. *StatPearls*. Treasure Island (FL) ineligible companies. Disclosure: Preeti Patel declares no relevant financial relationships with ineligible companies. Disclosure: Mayur Parmar declares no relevant financial relationships with ineligible companies.; 2024.

39. McCarthy MM, and Wright CL. Convergence of Sex Differences and the Neuroimmune System in Autism Spectrum Disorder. Biol Psychiatry. 2017;81(5):402-10.

40. Ribot J, Breton R, Calvo CF, Moulard J, Ezan P, Zapata J, et al. Astrocytes close the mouse critical period for visual plasticity. Science. 2021;373(6550):77-81.

41. Ackerman SD, Perez-Catalan NA, Freeman MR, and Doe CQ. Astrocytes close a motor circuit critical period. Nature. 2021;592(7854):414–20.

42. Banker SM, Gu X, Schiller D, and Foss-Feig JH. Hippocampal contributions to social and cognitive deficits in autism spectrum disorder. Trends Neurosci. 2021;44(10):793-807.

43. Roy DS, Zhang Y, Halassa MM, and Feng G. Thalamic subnetworks as units of function. Nat Neurosci. 2022;25(2):140-53.

44. Hwang BJ, Mohamed MA, and Brasic JR. Molecular imaging of autism spectrum disorder. Int Rev Psychiatry. 2017;29(6):530-54.

45. Perez-Garcia CG, and OʹLeary DDM. Formation of the Cortical Subventricular Zone Requires MDGA1-Mediated Aggregation of Basal Progenitors. Cell Rep. 2016;14(3):560-71.

46. Han KA, Kim YJ, Yoon TH, Kim H, Bae S, Um JW, et al. LAR-RPTPs Directly Interact with Neurexins to Coordinate Bidirectional Assembly of Molecular Machineries. J Neurosci. 2020;40(4):8438-62.

