## Supplementary Figures for "Bazedoxifene rescues sexually dimorphic autistic-like abnormalities in mice carrying a biallelic *MDGA1* mutation"

### Supplemental figures and figure legends

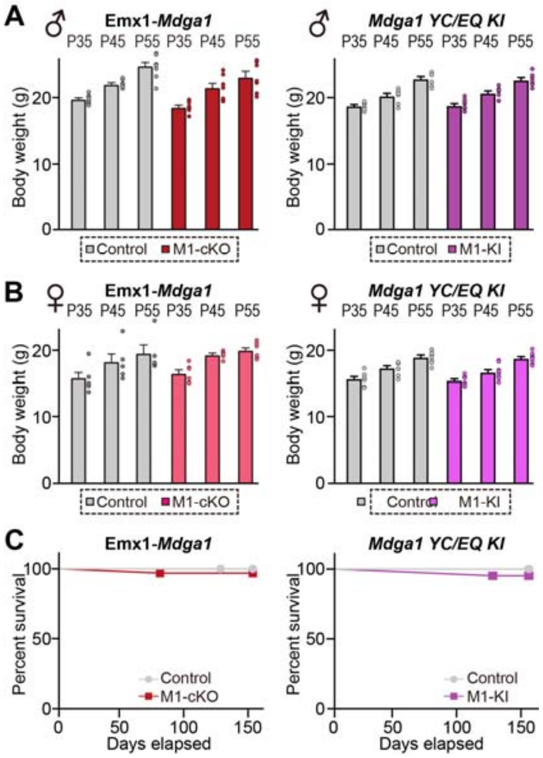


**Supplemental Figure 1. Analysis of body weight and the survival curve for *Mdga1*-cKO and *Mdga1*$^{Y636C/E751Q}$ KI mice.**

(**A**) Body weight measurements of male MDGA1 conditional knockout (cKO) (**left**) and MDGA1 Y636C/E751Q knock-in (KI) (**right**) mice compared to control mice. Body weight was recorded at postnatal days P35, P45, and P55. Data are presented as means ± SEMs (n = 7 mice/group).

(**B**) Body weight measurements of female MDGA1 conditional knockout (cKO) (**left**) and MDGA1 Y636C/E751Q KI (**right**) mice compared to control mice. Body weight was recorded at postnatal days P35, P45, and P55. Data are presented as means ± SEMs (n =5–9 mice/group).

(**C**) Survival curves showing the survival probability of male *Mdga1*-cKO and *Mdga1*$^{Y636C/E751Q}$ KI mice compared to control mice. The survival probability was tracked over time up to 150 days. Survival curves were generated using the Kaplan-Meier method (n = 20–32 mice/group).

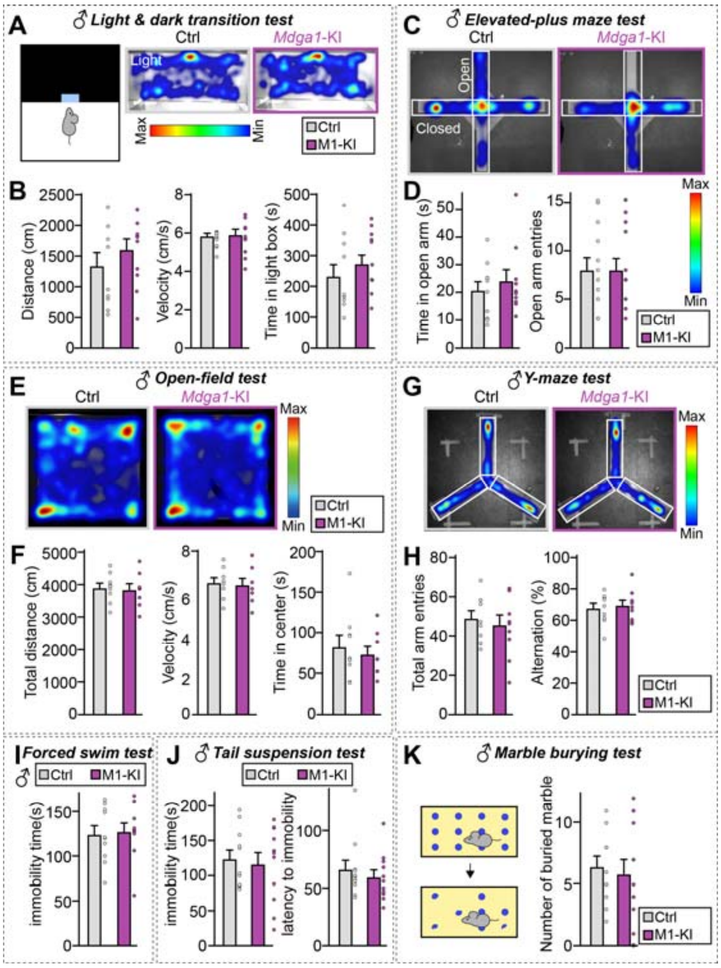


**Supplemental Figure 2. Analysis of behaviors of adult male *Mdga1*$^{Y636C/E751Q}$ KI mice.**

(**A** and **B**) Light and dark transition test results showing time spent in the light box (**A**), total distance traveled, and velocity for male control and *Mdga1*$^{Y636C/E751Q}$ KI mice (**B**). Data are presented as means ± SEMs (n = 9–10 mice/group).

(**C** and **D**) Elevated-plus maze test results showing time spent in the open arms (**C**) and number of open arm entries for male control and *Mdga1*$^{Y636C/E751Q}$ KI mice (**D**). Data are presented as means ± SEMs (n = 10 mice/group).

(**E** and **F**) Open-field test results showing total distance traveled (**E**), time spent in the center, and velocity for male control and *Mdga1*$^{Y636C/E751Q}$ KI mice (**F**). Data are presented as means ± SEMs (n = 8–9 mice/group).

(**G** and **H**) Y-maze test results showing total arm entries (**G**) and alternation percentage for male control and *Mdga1*$^{Y636C/E751Q}$ KI mice (**H**). Data are presented as means ± SEMs (n = 9–10 mice/group).

(**I**) Forced swim test results showing total immobility time for male control and *Mdga1*$^{Y636C/E751Q}$ KI mice. Data are presented as means ± SEMs (n = 7 mice/group).

(**J**) Tail suspension test results showing the latency to immobility and total immobility time for male control and *Mdga1*$^{Y636C/E751Q}$ KI mice. Data are presented as means ± SEMs (n = 7 mice/group).

(**K**) Marble burying test results showing the number of marbles buried by male control and *Mdga1*$^{Y636C/E751Q}$ KI mice. Data are presented as means ± SEMs (n = 11–13 mice/group.)

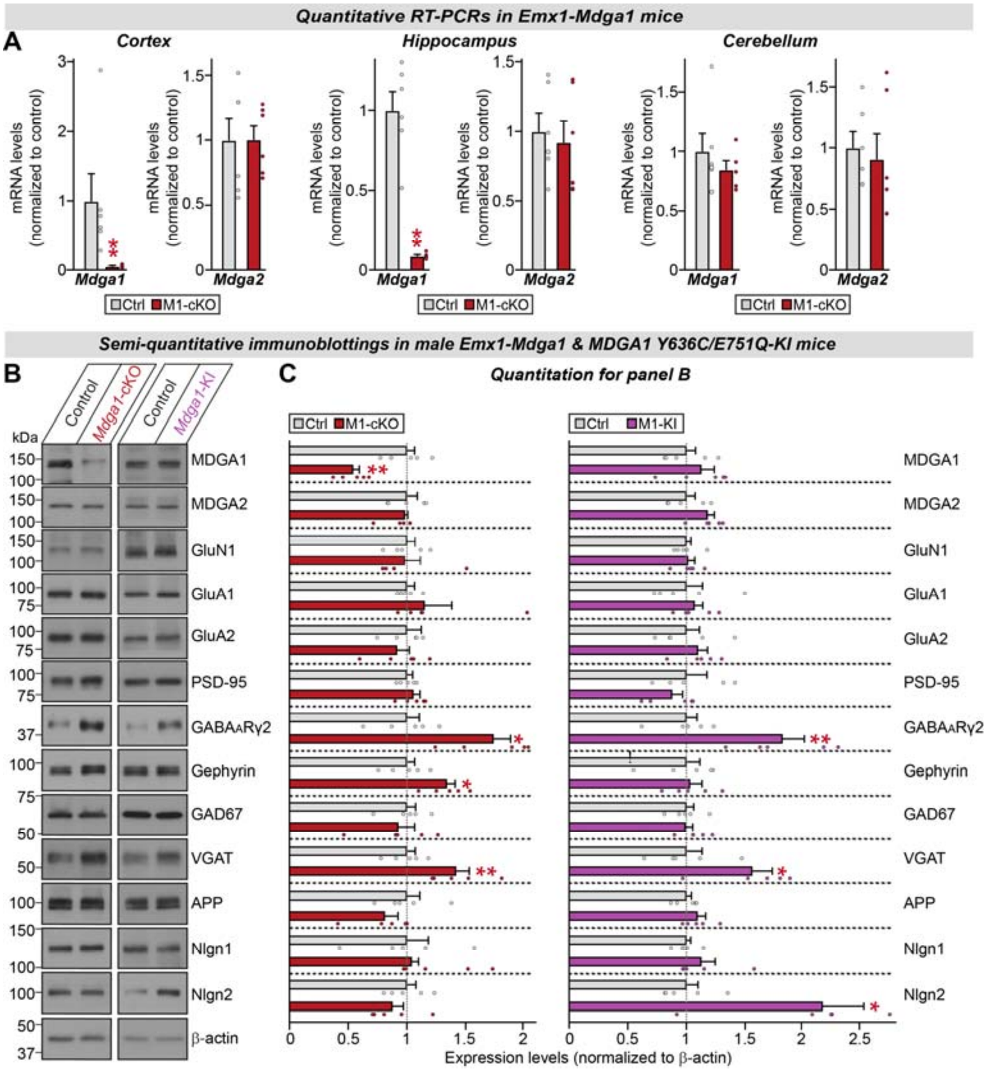


**Supplemental Figure 3. Analysis of expression levels of *Mdga* mRNAs and MDGA proteins as well as expression levels of other synaptic proteins in the hippocampus of male *Mdga1*-cKO and male *Mdga1*$^{Y636C/E751Q}$ KI mice.**

(**A**) Quantitative RT-PCR analysis of *Mdga1* and *Mdga2* mRNA levels in the hippocampus, cortex, and cerebellum of adult male *Mdga1*-cKO mice. Data are normalized to control levels and presented as means ± SEMs (n = 5–6 mice/group; $^{**}p < 0.01$, Mann–Whitney *U* test).

(**B**) Semi-quantitative immunoblotting of various synaptic proteins in the hippocampus of adult male control, *Mdga1*-cKO and *Mdga1*$^{Y636C/E751Q}$ KI mice. β-actin was used as a loading control for normalization.

(**C**) Semi-quantitative analysis of immunoblotting results shown in panel (**B**). Expression levels of various proteins are normalized to β-actin and presented as fold changes relative to control mice. Data are presented as means ± SEMs (n = 5 mice/group; $^{*}p < 0.05$, $^{**}p < 0.01$; Mann–Whitney *U* test).

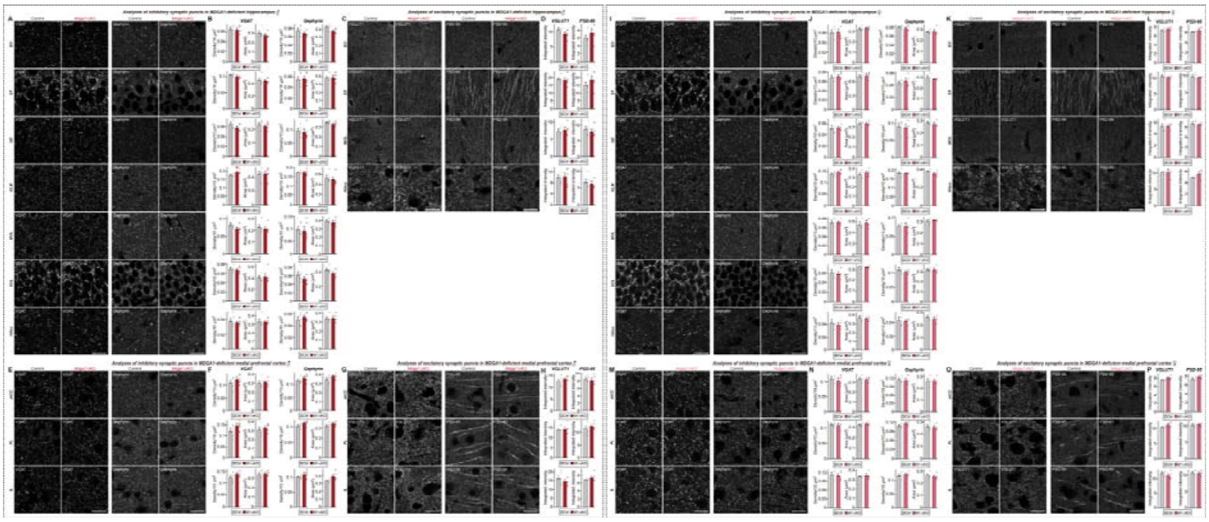

**Supplemental Figure 4. Analysis of glutamatergic and GABAergic synaptic puncta in the hippocampus and mPFC of adult *Mdga1*-cKO mice.**

(**A**) Representative images of GABAergic synaptic puncta in the hippocampus of male control and *Mdga1*-cKO mice. Neurons were immunostained for VGAT and gephyrin in various hippocampal layers. Abbreviation: SO, *stratum oriens*; SR, *stratum radiatum*; SLM, *stratum lacunosum moleculare*; SP, *stratum pyramidale*; MOL, molecular layer; and GCL, granule cell layer. Scale bar, 10 μm (applies to all images).

(**B**) Quantification of GABAergic synaptic puncta density and area in the hippocampus of male control and *Mdga1*-cKO mice. Data are presented as means ± SEMs (n = 5 mice/group; $*p < 0.05$; Mann–Whitney *U* test).

(**C**) Representative images of glutamatergic synaptic puncta in the hippocampus of male control and *Mdga1*-cKO mice. Neurons were immunostained for VGLUT1 and PSD-95 in various hippocampal layers. Scale bar, 10 μm (applies to all images).

(**D**) Quantification of glutamatergic synaptic puncta density and area in the hippocampus of male control and *Mdga1*-cKO mice. Data are presented as means ± SEMs (n = 5 mice/group).

(**E**) Representative images of GABAergic synaptic puncta in the mPFC of male control and *Mdga1*-cKO mice. Neurons were immunostained for VGAT and gephyrin in different layers of the mPFC. Abbreviations: ACC, anterior cingulate cortex; IL, infralimbic cortex; PL, prelimbic cortex. Scale bar, 10 μm (applies to all images).

(**F**) Quantification of GABAergic synaptic puncta density and area in the mPFC of male control and *Mdga1*-cKO mice. Data are presented as means ± SEMs (n = 5 mice/group).

(**G**) Representative images of glutamatergic synaptic puncta in the mPFC of male control and *Mdga1*-cKO mice. Neurons were immunostained for VGLUT1 and PSD-95 in different layers of the mPFC. Scale bar, 10 μm (applies to all images).

(**H**) Quantification of glutamatergic synaptic puncta density and area in the mPFC of male control and *Mdga1*-cKO mice. Data are presented as means ± SEMs (n = 5 mice/group).

(**I**) Representative images of GABAergic synaptic puncta in the hippocampus of female control and *Mdga1*-cKO mice. Scale bar, 10 μm (applies to all images).

(**J**) Quantification of GABAergic synaptic puncta density and area in the hippocampus of female control and *Mdga1*-cKO mice. Data are presented as means ± SEMs (n = 4 mice/group).

(**K**) Representative images of glutamatergic synaptic puncta in the hippocampus of female control and *Mdga1*-cKO mice. Neurons were immunostained for VGLUT1 and PSD-95 in various hippocampal layers. Scale bar, 10 μm (applies to all images).

(**L**) Quantification of glutamatergic synaptic puncta density and area in the hippocampus of female control and *Mdga1*-cKO mice. Data are presented as means ± SEMs (n = 4 mice/group).

(**M**) Representative images of GABAergic synaptic puncta in the mPFC of female control and *Mdga1*-cKO mice. Neurons were immunostained for VGAT and gephyrin in different layers of the mPFC. Scale bar, 10 µm (applies to all images).
(**N**) Quantification of GABAergic synaptic puncta density and area in the mPFC of female control and *Mdga1*-cKO mice. Data are presented as means ± SEMs (n = 4 mice/group).
(**O**) Representative images of glutamatergic synaptic puncta in the mPFC of female control and *Mdga1*-cKO mice. Neurons were immunostained for VGLUT1 and PSD-95 in different layers of the mPFC. Scale bar, 10 µm (applies to all images).
(**P**) Quantification of glutamatergic synaptic puncta density and area in the mPFC of female control and *Mdga1*-cKO mice. Data are presented as means ± SEMs (n = 4 mice/group).

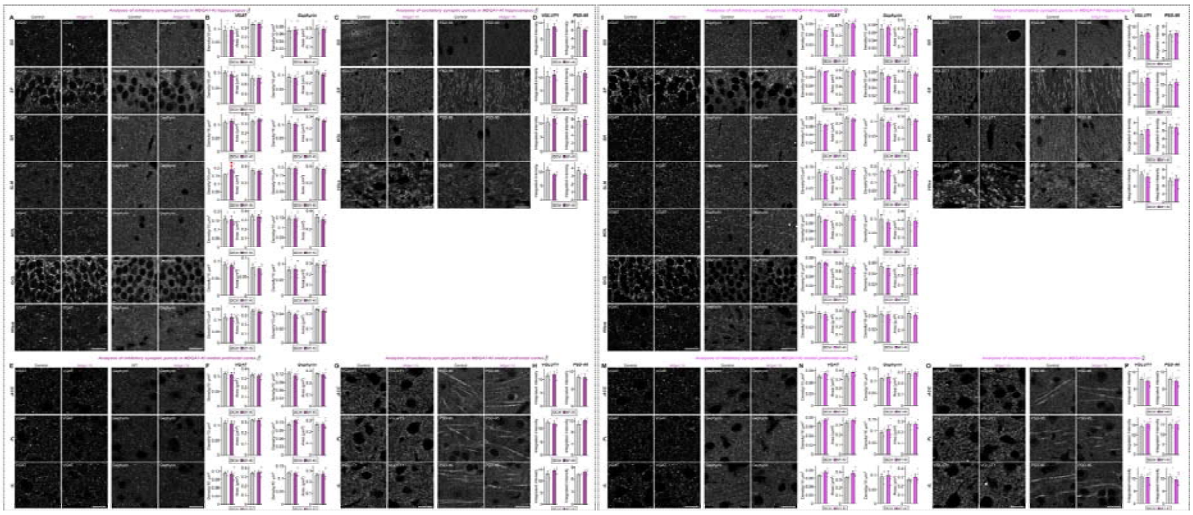

**Supplemental Figure 5. Analysis of glutamatergic and GABAergic synapti puncta in the hippocampus and mPFC of adult *Mdga1*$^{Y636C/E751Q}$ KI mice.**

(**A**) Representative images of GABAergic synaptic puncta in the hippocampus of male control and *Mdga1*$^{Y636C/E751Q}$ KI mice. Neurons were immunostained for VGAT and gephyrin in various hippocampal layers. Scale bar, 10 μm (applies to all images).

(**B**) Quantification of GABAergic synaptic puncta density and area in the hippocampus of male control and *Mdga1*$^{Y636C/E751Q}$ KI mice. Data are presented as means ± SEMs (n = 5 mice/group; $^{**}p < 0.01$; Mann–Whitney *U* test).

(**C**) Representative images of glutamatergic synaptic puncta in the hippocampus of male control and *Mdga1*$^{Y636C/E751Q}$ KI mice. Neurons were immunostained for VGLUT1 and PSD-95 in various hippocampal layers. Scale bar, 10 μm (applies to all images).

(**D**) Quantification of glutamatergic synaptic puncta density and area in the hippocampus of male control and *Mdga1*$^{Y636C/E751Q}$ KI mice. Data are presented as means ± SEMs (n = 5 mice/group).

(**E**) Representative images of GABAergic synaptic puncta in the mPFC of male control and *Mdga1*$^{Y636C/E751Q}$ KI mice. Neurons were immunostained for VGAT and gephyrin in different layers of the mPFC. Scale bar: 10 μm (applies to all images).

(**F**) Quantification of GABAergic synaptic puncta density and area in the mPFC of male control and *Mdga1*$^{Y636C/E751Q}$ KI mice. Data are presented as means ± SEMs (n = 5 mice/group).

(**G**) Representative images of glutamatergic synaptic puncta in the mPFC of male control and *Mdga1*$^{Y636C/E751Q}$ KI mice. Neurons were immunostained for VGLUT1 and PSD-95 in different layers of the mPFC. Scale bar, 10 μm (applies to all images).

(**H**) Quantification of glutamatergic synaptic puncta density and area in the mPFC of male control and *Mdga1*$^{Y636C/E751Q}$ KI mice. Data are presented as means ± SEMs (n = 5 mice/group).

(**I**) Representative images of GABAergic synaptic puncta in the hippocampus of female control and *Mdga1*$^{Y636C/E751Q}$ KI mice. Neurons were immunostained for VGAT and gephyrin in various hippocampal layers. Scale bar, 10 μm (applies to all images).

(**J**) Quantification of GABAergic synaptic puncta density and area in the hippocampus of female control and *Mdga1*$^{Y636C/E751Q}$ KI mice. Data are presented as means ± SEMs (n = 5 mice/group).

(**K**) Representative images of glutamatergic synaptic puncta in the hippocampus of female control and *Mdga1*$^{Y636C/E751Q}$ KI mice. Neurons were immunostained for VGLUT1 and PSD-95 in various hippocampal layers. Scale bar, 10 μm (applies to all images).

(**L**) Quantification of glutamatergic synaptic puncta density and area in the hippocampus of female control and *Mdga1*$^{Y636C/E751Q}$ KI mice. Data are presented as means ± SEMs (n = 5 mice/group).

(**M**) Representative images of GABAergic synaptic puncta in the mPFC of female control and *Mdga1*$^{Y636C/E751Q}$ KI mice. Neurons were immunostained for VGAT and gephyrin in different layers of the mPFC. Scale bar, 10 μm (applies to all images).

(**N**) Quantification of GABAergic synaptic puncta density and area in the mPFC of female control and *Mdga1*$^{Y636C/E751Q}$ KI mice. Data are presented as means ± SEMs (n = 5 mice/group).
(**O**) Representative images of excitatory synaptic puncta in the mPFC of female control and *Mdga1*$^{Y636C/E751Q}$ KI mice. Neurons were immunostained for VGLUT1 and PSD-95 in different layers of the mPFC. Scale bar, 10 μm (applies to all images).
(**P**) Quantification of glutamatergic synaptic puncta density and area in the mPFC of female control and *Mdga1*$^{Y636C/E751Q}$ KI mice. Data are presented as means ± SEMs (n = 5 mice/group).

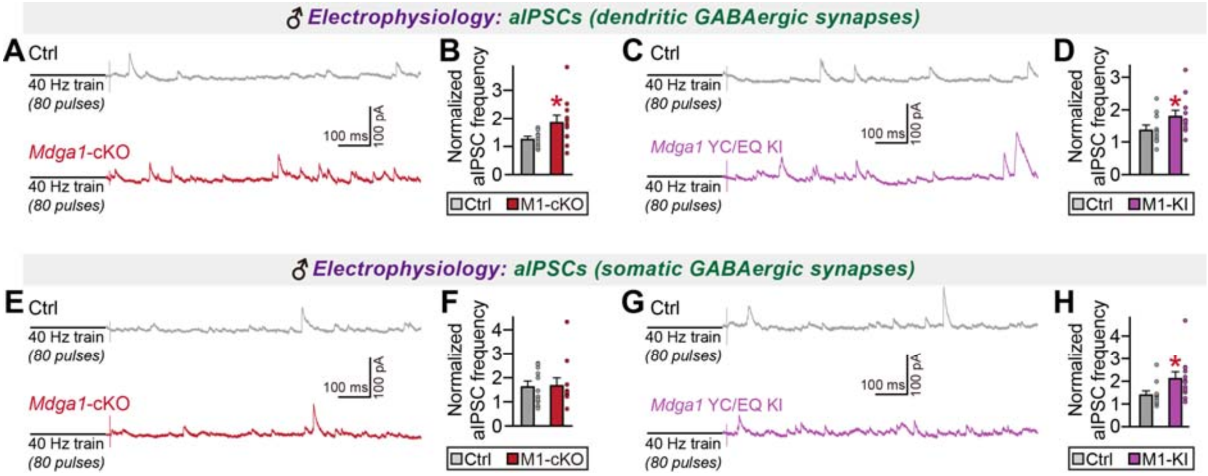


**Supplemental Figure 6. Analysis of asynchronous GABAergic evoked synaptic transmission in the hippocampal CA1 pyramidal neurons of adult male *Mdga1*-cKO and *Mdga1*$^{\mathbf{Y636C/E751Q}}$ KI mice.**

(**A** and **B**) Representative traces (**A**) and quantification of normalized asynchronous inhibitory postsynaptic currents (aIPSCs) frequency (**B**) recorded from dendritic GABAergic synapses in male control and *Mdga1*-cKO mice. Data are presented as means ± SEMs (n = 11 cells/group; **p* < 0.05; Mann–Whitney *U* test).

(**C** and **D**) Representative traces (**C**) and quantification of normalized aIPSC frequency (**D**) in dendritic GABAergic synapses in male control and *Mdga1*$^{Y636C/E751Q}$ KI mice. Data are presented as means ± SEMs (n = 11–12 cells/group; **p* < 0.05; Mann–Whitney *U* test).

(**E** and **F**) Representative traces (**E**) and quantification of normalized aIPSCs frequency (**F**) recorded from somatic GABAergic synapses in male control and *Mdga1*-cKO mice. Data are presented as means ± SEMs (n = 11 cells/group).

(**G** and **H**) Representative traces (**G**) and quantification of normalized aIPSC frequency (**H**) in somatic GABAergic synapses in male control and *Mdga1*$^{Y636C/E751Q}$ KI mice. Data are presented as means ± SEM (n = 11–12 cells/group; **p* < 0.05; Mann–Whitney *U* test).

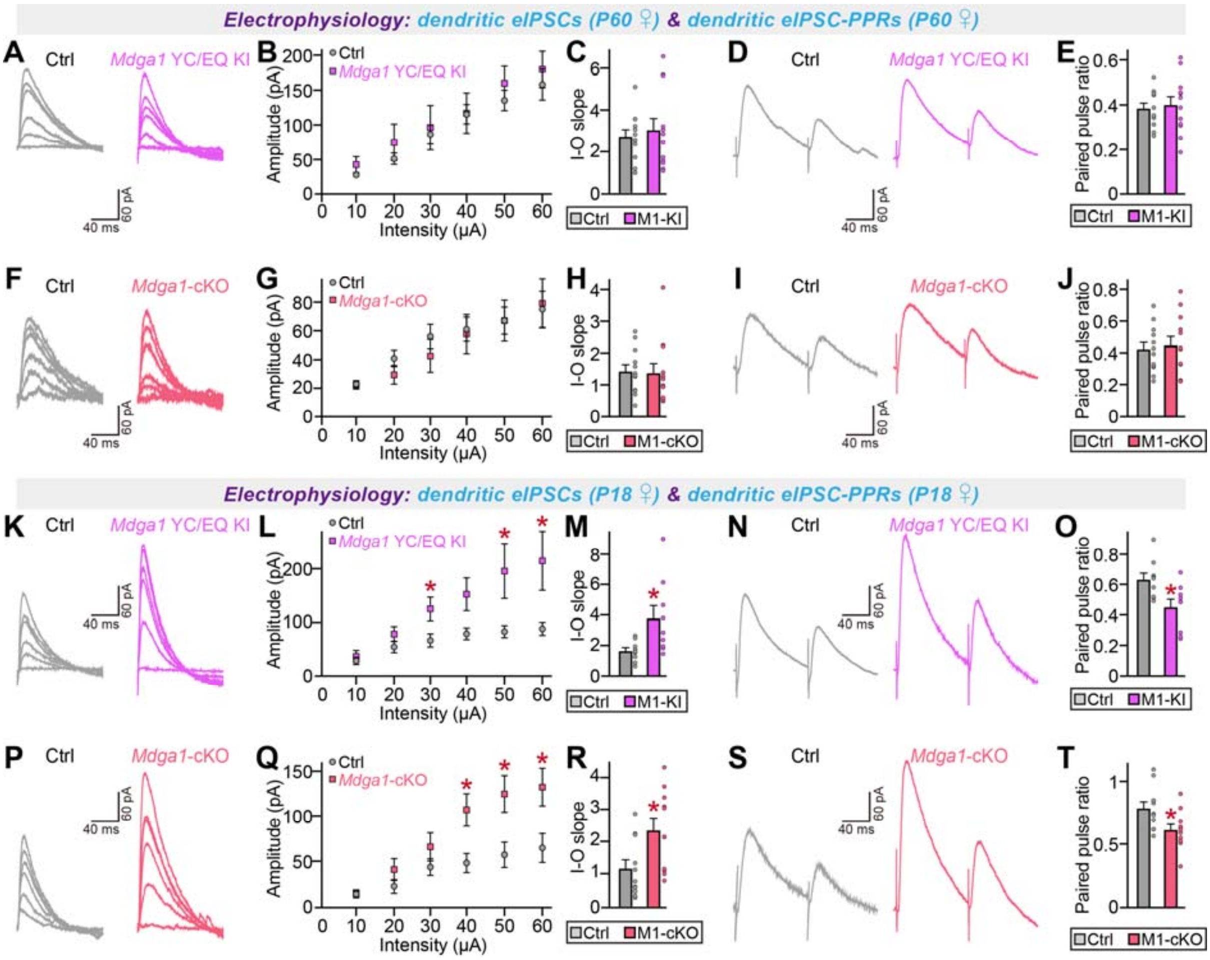


**Supplemental Figure 7. Analysis of synchronous GABAergic evoked synaptic transmission in the hippocampal CA1 pyramidal neurons of adult female *Mdga1*-cKO and *Mdga1*$^{Y636C/E751Q}$ KI mice.**

(**A–E**) Representative traces (**A**), input-output (I-O) curves (**B** and **C**), and paired-pulse ratios (PPRs) (**D** and **E**) of dendritic eIPSCs recorded from CA1 pyramidal neurons in adult female control and *Mdga1*$^{Y636C/E751Q}$ KI mice. Data are presented as means ± SEMs (n = 12 cells/group).

(**F–J**) Representative traces (**F**), I-O curves (**G** and **H**), and PPRs (**I** and **J**) of dendritic eIPSCs recorded from CA1 pyramidal neurons in adult female control and *Mdga1*-cKO mice. Data are presented as means ± SEMs (n = 12 cells/group).

(**K–O**) Representative traces (**K**), I-O curves (**L** and **M**), and PPRs (**N** and **O**) of dendritic eIPSCs recorded from CA1 pyramidal neurons in P18 female control and *Mdga1*$^{Y636C/E751Q}$ KI mice. Data are presented as means ± SEMs (n = 9 cells/group; *$p < 0.05$; Mann–Whitney *U* test).

(**P–T**) Representative traces (**P**), I-O curves (**Q** and **R**), and PPRs (**S** and **T**) of dendritic eIPSCs recorded from CA1 pyramidal neurons in P18 female control and *Mdga1*-cKO mice. Data are presented as means ± SEMs (n = 11 cells/group; *$p < 0.05$; Mann–Whitney *U* test).

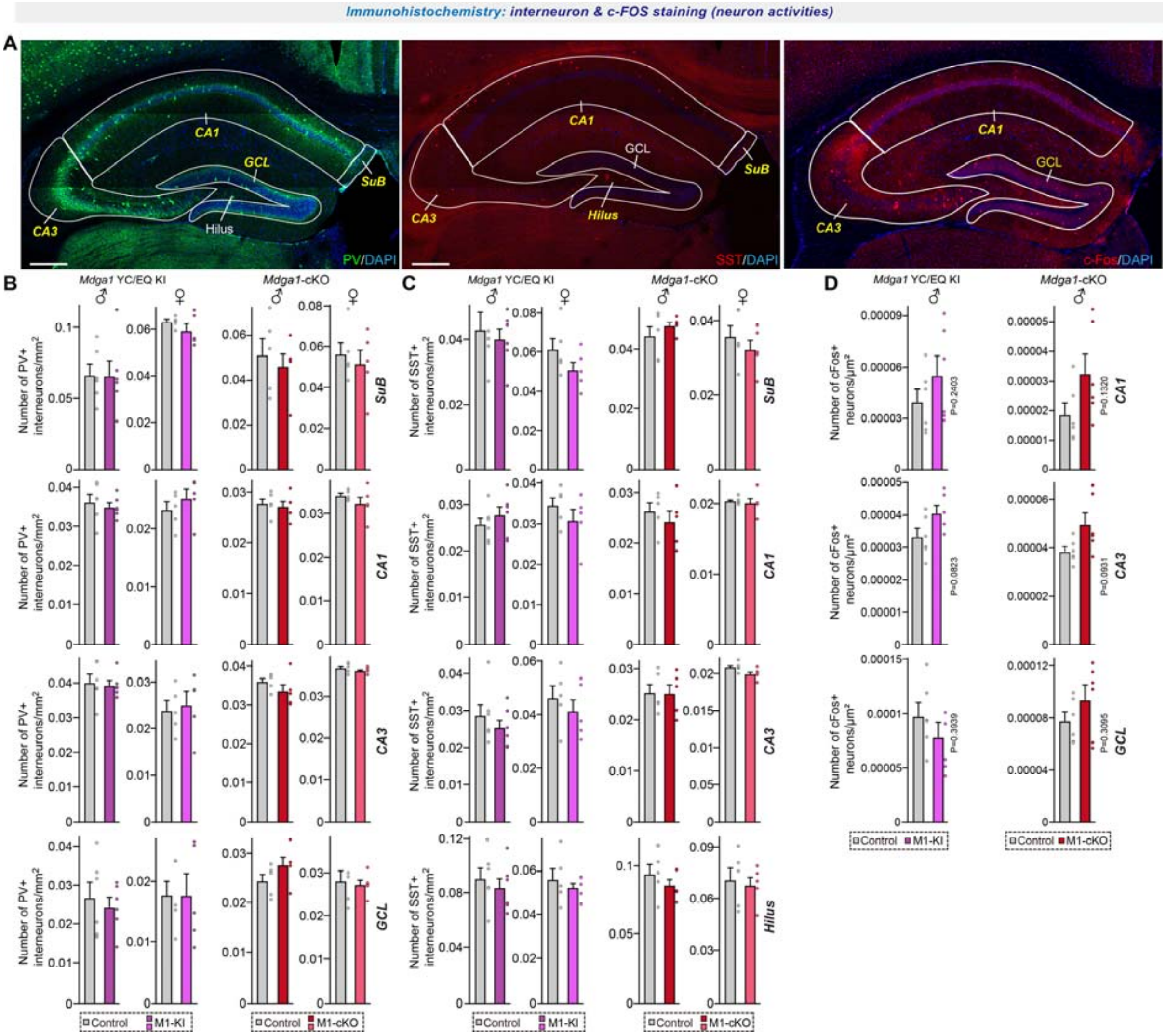


**Supplemental Figure 8. Analysis of density of interneurons and c-FOS+ neurons in the hippocampus of adult *Mdga1*-cKO and *Mdga1*$^{Y636C/E751Q}$ KI mice.**

(**A**) Representative images of parvalbumin (PV)$^{+}$ (**left**), somatostatin (SST)$^{+}$ (**middle**), and c-FOS+ neurons (**right**) in various hippocampal layers of control, *Mdga1*-cKO and *Mdga1*$^{Y636C/E751Q}$ KI mice. Scale bar, 200 μm (applies to all images).

(**B**) Quantification of PV$^{+}$ interneuron density in the hippocampal layer of control, *Mdga1*-cKO and *Mdga1*$^{Y636C/E751Q}$ KI mice, separated by sex (male and female). Data are presented as means ± SEMs (n = 5–6 mice/group). Abbreviations: SuB, subiculum; GCL, granular cell layer.

(**C**) Quantification of SST$^{+}$ interneuron density in hippocampal layers of control, *Mdga1*-cKO and *Mdga1*$^{Y636C/E751Q}$ KI mice, separated by sex (male and female). Data are presented as means ± SEMs (n = 5–6 mice/group).

(**D**) Quantification of c-FOS$^{+}$ neuron density (c-FOS/DAPI) in various hippocampal layers of male control, *Mdga1*-cKO and *Mdga1*$^{Y636C/E751Q}$ KI mice. Data are presented as means ± SEMs (n = 6 mice/group).

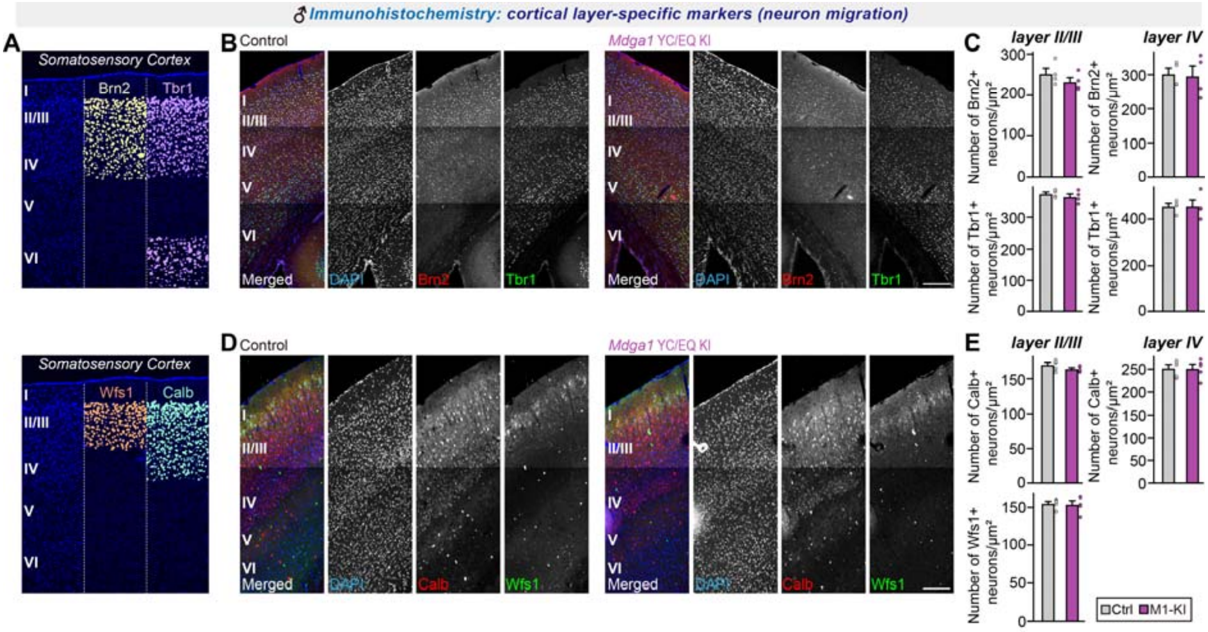


**Supplemental Figure 9. Analysis of cortical neuron migration for adult male *Mdga1*$^{\mathrm{Y636C/E751Q}}$ KI mice.**

(**A**) Schematic of cortical layer-specific markers used in the current study.

(**B**) Representative images of Brn2- and Tbr1-labeled neurons in the somatosensory cortex of adult male control and *Mdga1*$^{\mathrm{Y636C/E751Q}}$ KI mice. Neurons were immunostained with antibodies against Brn2 (red), Tbr1 (green) and DAPI (blue). Brn2 and Tbr1 are cortical layer-specific markers (Brn2 for layers II–IV; Tbr1 for layers II/III and primarily VI). Scale bar, 200 μm (applies to all images).

(**C**) Quantification of Brn2$^{+}$ and Tbr1$^{+}$ neuron density in cortical layers (layers II/III and IV) of adult male control and *Mdga1*$^{\mathrm{Y636C/E751Q}}$ KI mice. Data are presented as means ± SEMs (n = 5 mice/group; Mann–Whitney *U* test).

(**D**) Representative images of calbindin- and Wfs1-labeled neurons in the somatosensory cortex of adult male control and *Mdga1*$^{\mathrm{Y636C/E751Q}}$ KI mice. Neurons were immunostained with antibodies against calbindin (Calb; red), Wfs1 (green) and DAPI (blue). Calb and Wfs1 are cortical layer-specific markers (Calb for layers II–IV; Wfs1 for layers II/III). Scale bar, 200 μm (applies to all images).

(**E**) Quantification of calbindin$^{+}$ and Wfs1$^{+}$ neuron density in cortical layers II/III and IV of adult male control and *Mdga1*$^{\mathrm{Y636C/E751Q}}$ KI mice. Data are presented as means ± SEMs (n = 5 mice/group; Mann–Whitney *U* test).

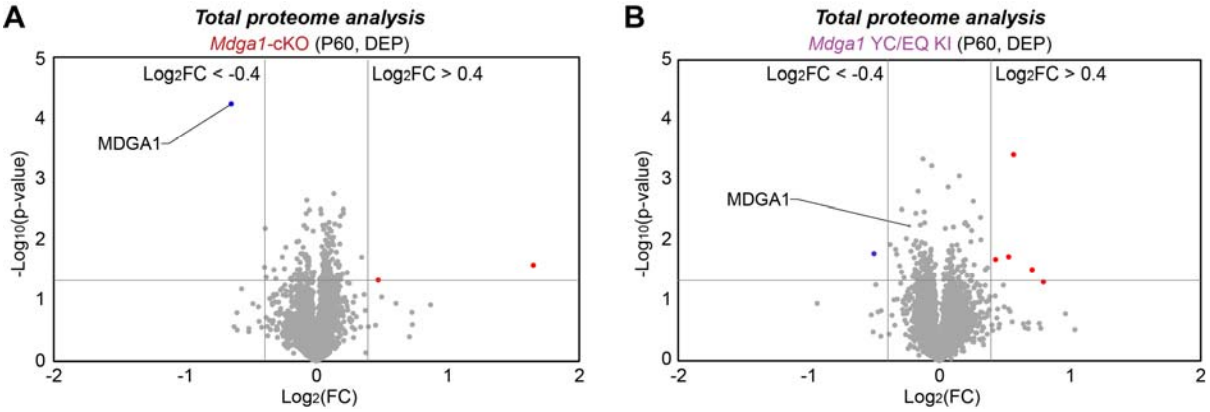


**Supplemental Figure 10. Volcano plot of proteins identified from proteomic analyses of hippocampi from adult *Mdga1*-cKO and *Mdga1*$^{Y636C/E751Q}$ KI mice.**

(**A**) Volcano plot displaying differentially expressed proteins (DEPs) identified from hippocampi of adult male *Mdga1*-cKO mice compared to controls. Proteins with $Log_2$ fold change ($Log_2FC$) < -0.4 are indicated in blue, while those with $Log_2FC$ > 0.4 are shown in red. MDGA1 is highlighted among the few downregulated proteins.

(**B**) Same as (**A**), except showing results obtained from adult male *Mdga1*$^{Y636C/E751Q}$ KI mice compared to controls. Each dot represents an individual protein, with significance represented by $-Log_{10}$(*p*-value) on the y-axis and $Log_2FC$ on the x-axis.

A

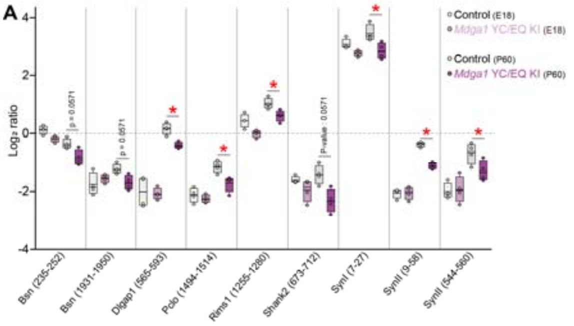


B

| Protein name | Sequence |
|---|---|
| Bsn | (235-252) SQQQLHSPALSPAHSPAK; 2 phosphorylation |
| Bsn | (1931-1950) SVTDTALPGQSSGPFYSPR; 1 phosphorylation |
| Dlgap1 | (565-593) TTTKPFISITAQSSTESAQDAYMDGQGQR; 1 phosphorylation |
| Pclo | (1494-1514) QIIEMSADEDASGSEDEEFIR; 1 phosphorylation |
| Rims1 | (1255-1280) QPSRESTDGSINSYSSEGNLIFPGVR; 1 phosphorylation |
| Shank2 | (673-712) SSQGSSMEIDPQATEPGQLRPDDSLTVSSPFAAAIAGAVR; 1 phosphorylation |
| SynI | (7-27) RLSDSNFMANLPNGYMDLQR; 1 phosphorylation |
| SynII | (9-58) LSDSSFIANLPNGYMTDLQRPEPQQPPPAPGPGAATASAATSAASPGPER; 1 phosphorylation |
| SynII | (544-560) SQSLTNAFSFSESSFFR; 1 phosphorylation |

**Supplemental Figure 11. Analysis of phosphorylation sites that are altered in adult male *Mdga1*$^{Y636C/E751Q}$ KI mice.**

(**A**) $Log_2$ ratio of phosphopeptides representing the proteins, SynI, SynII, Piccolo (Pclo), Bassoon (Bsn), Shank2, RIMS1, and Dlgap1, in embryos (E18) and adult (P60) male *Mdga1*$^{Y636C/E751Q}$ KI mice, compared to controls. Data are presented as means ± SEMs (n = 4 mice/group; *$p < 0.05$; two-way ANOVA followed by Tukey's *post-hoc* test).

(**B**) A table listing the identified phosphopeptides and their corresponding proteins.

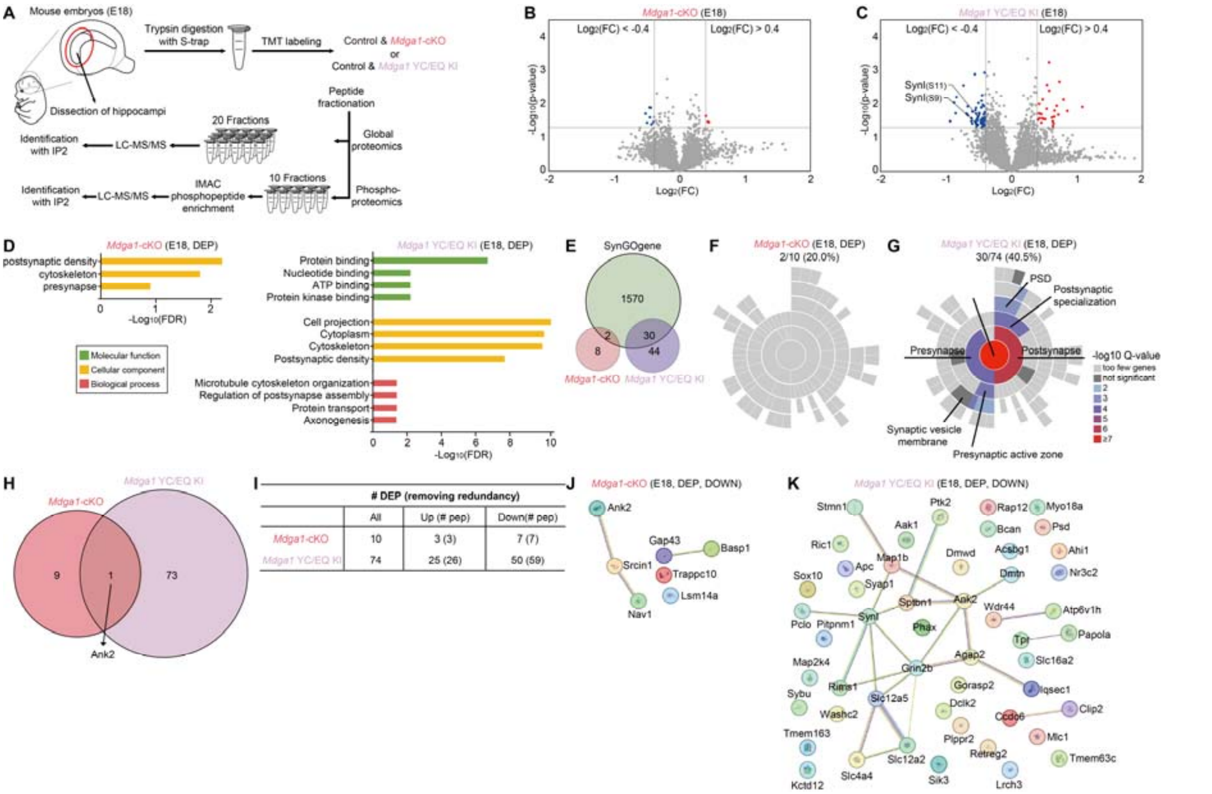


### Supplemental Figure 12. Phosphoproteomic analysis of hippocampi from embryos of *Mdga1*-cKO and *Mdga1*$^{Y636C/E751Q}$ KI mice.

(**A**) Schematic diagram of phosphoproteomic analysis of hippocampal lysates from embryos of *Mdga1*-cKO, *Mdga1*$^{Y636C/E751Q}$ KI, and littermate control mice.

(**B**) Volcano plot of 4,338 phosphopeptides identified from hippocampal lysates of *Mdga1*-cKO embryos. Differentially expressed phosphopeptides (DEPPs) with significant increases or decreases (Log$_2$FC < -0.4 or > 0.4; $p < 0.05$) are shown in red (3 phosphopeptides) and blue (7 phosphopeptides), respectively.

(**C**) Volcano plot of 5,124 phosphopeptides identified from hippocampal lysates of *Mdga1*$^{Y636C/E751Q}$ KI embryos. DEPPs with significant increases or decreases (Log$_2$FC < -0.4 or > 0.4; $p < 0.05$) are shown in red (26 phosphopeptides) and blue (59 phosphopeptides), respectively.

(**D**) Gene ontology (GO) analysis was performed on differentially expressed phosphoproteins (DEPs) identified from the phosphoproteomic analyses of *Mdga1*-cKO (**left**) and *Mdga1*$^{Y636C/E751Q}$ KI (**right**) embryos. The bar charts depict the significantly enriched GO terms in the Molecular Function (green), Cellular Component (yellow), and Biological Process (red) categories, represented as -log$_{10}$(FDR) values.

(**E**) Ven diagram visualization of SynGO category proteins.

(**F**) Localization annotations of SynGO category proteins from *Mdga1*-cKO embryos.

(**G**) Localization annotations of SynGO category proteins from *Mdga1*$^{Y636C/E751Q}$ KI embryos.

(**H**) Venn diagram showing the overlap of DEPs between *Mdga1*-cKO and *Mdga1*$^{Y636C/E751Q}$ KI embryos. Ank2 is shared between the conditions; 9 DEPs are unique to *Mdga1*-cKO embryos and 73 are unique to *Mdga1*$^{Y636C/E751Q}$ KI embryos.

(**I**) Summary table of DEPs after removal of redundancy. The table shows the total number DEPs with a breakdown of the upregulated and downregulated proteins in *Mdga1*-cKO and *Mdga1*$^{Y636C/E751Q}$ KI embryos, together with the number of unique peptides (# pep).

(**J**) STRING analysis of downregulated DEPs from *Mdga1*-cKO embryos ($p < 0.05$ and Log$_2$FC < -0.4).

(**K**) STRING analysis of downregulated DEPs from *Mdga1*$^{Y636C/E751Q}$ KI embryos ($p < 0.05$ and Log$_2$FC < -0.4).

**Supplemental Table 1. List of total proteins used for volcano plot analysis of results from proteomic analysis of hippocampal tissues from control and *Mdga1*-cKO adult mice and embryos.**

**Supplemental Table 2. List of total proteins used for volcano plot analysis of results from proteomic analysis of hippocampal tissues from control and *Mdga1*$^{Y636C/E751Q}$ KI adult mice and embryos.**

**Supplemental Table 3. List of phosphopeptides used for volcano plot analysis of results from phosphoproteomic analysis of hippocampal tissues from control and *Mdga1*-cKO adult mice and embryos.**

**Supplemental Table 4. List of phosphopeptides used for volcano plot analysis of results from proteomic analysis of hippocampal tissues from control and *Mdga1*$^{Y636C/E751Q}$ KI adult mice and embryos.**

**Supplemental Table 5. Differentially expressed phosphopeptides (DEPPs) exhibiting significant ($p$ < 0.05 & $log_2FC$ < -0.4 or > 0.4) differences between adult *Mdga1*-cKO and *Mdga1*$^{Y636C/E751Q}$ KI mice versus adult control mice.**

**Supplemental Table 6. Differentially expressed phosphopeptides (DEPPs) exhibiting significant ($p$ < 0.05 & $log_2FC$ < -0.4 or > 0.4) differences between *Mdga1*-cKO embryos and *Mdga1*$^{Y636C/E751Q}$ KI embryos versus control embryos.**
